# Stress, Epigenetic Remodeling and FKBP51: Pathways to Chronic Pain Vulnerability

**DOI:** 10.1101/2025.01.07.631709

**Authors:** Oakley B. Morgan, Samuel Singleton, Sara Hestehave, Tim Sarter, Eva Wozniak, Charles A Mein, Felix Hausch, Christopher G. Bell, Sandrine M. Géranton

**Affiliations:** Dept. of Cell & Developmental Biology, University College London, WC1E 6BT, United Kingdom; School of Medicine, University of Dundee, Dundee, DD1 5EH, United Kingdom; Dept. Of Experimental Medicine, University of Copenhagen, Copenhagen N 2200, Denmark; Department of Pharmacy, Pharmaceutical Technology and Biopharmaceutics, Ludwig-Maximilians-Universität München, Munich, 81377, Germany; Genome Centre, Faculty of Medicine and Dentistry, Queen Mary University of London, London, E1 2AT, UK; Institute of Organic Chemistry and Biochemistry, Technical University Darmstadt, Darmstadt, 64287, Germany; William Harvey Research Institute, Barts & The London Faculty of Medicine, Charterhouse Square, Queen Mary University of London, London, EC1M 6BQ, United Kingdom; QMUL Centre for Epigenetics, Queen Mary University of London, London, E1 4NS, United Kingdom

**Keywords:** Stress, chronic pain, epigenetics, DNA methylation, FKBP51

## Abstract

Stress is thought to contribute to the persistence of pain and comorbid anxiety, yet the underlying mechanisms remain unclear. In our pre-clinical model, sub-chronic stress exacerbated subsequently induced inflammatory pain and accelerated the development of comorbid anxiety. DNA methylation analysis of spinal cord tissue after stress exposure revealed hypomethylation in the *Fkbp5* promoter site for the canonical FKBP51 transcript and other stress-related genes. However, most epigenetic changes in key regulatory regions did not correlate with changes in gene expression assessed by RNA sequencing, suggesting that stress exposure had remodeled the epigenome without altering gene activity and primed genes for hyper-responsiveness to future challenges. FKBP51 inhibition during stress exposure reduced the exacerbation of inflammatory pain by stress and reversed several stress-induced DNA methylation changes in promoter regions of genes associated with stress and nociception, including *Rtn4, Cdk5* and *Nrxn1*, but not *Fkbp5*. These results indicate that sub-chronic stress leads to the hypomethylation of *Fkbp5* and increased susceptibility to chronic pain driven by FKBP51, but reversing *Fkbp5* hypomethylation is not necessary to prevent chronic pain vulnerability, which is likely driven by complex epigenetic regulation of multiple stress-regulated genes.

## 1. Introduction

Chronic stress is a well-known modulator of physiological and behavioral responses, often leading to increased vulnerability to various pathological conditions^1^. Prolonged exposure to stress activates the hypothalamic-pituitary-adrenal (HPA) axis, leading to elevated levels of glucocorticoids, which can disrupt normal bodily functions^2^. This dysregulation affects immune responses, promotes inflammation and alters neural circuits, particularly in regions associated with mood. Consequently, chronic stress is linked to a heightened risk of developing disorders such as anxiety, depression, cardiovascular diseases and chronic pain^3–5^. Stressful experiences indeed have a profound impact on the manifestation of pain, both in humans and rodents. While short-lasting, intense stress experiences can trigger a physiological reaction that attenuates pain signalling and allows for expression of protective behaviours, sustained exposure to stress exacerbates hyperalgesic states^6, 7^. Moreover, stress experience can prime for hyper-responsiveness to subsequent injury, as a result of long-lasting changes involving complex peripheral, central and systemic mechanisms^8^ and stressful life events are considered a key environmental factor in increased risk for developing chronic pain^9, 10^.

FK 506 binding protein 51 (FKBP51) is a crucial element of the stress axis that regulates the glucocorticoid receptor (GR) sensitivity to glucocorticoids^11, 12^. and *FKBP5* genetic polymorphisms and DNA methylation landscape have both been associated with increased risk for developing mental health disorders in small-scale human studies^13–16^. This includes the observation of early life adversity in humans reducing *FKBP5* DNA methylation (DNAm) at the promoter region, an epigenetic change that influences the gene’s expression, its responsiveness to stress and susceptibility to psychiatric disorders^14, 17^. While these genetic findings have not, to date, been replicated in larger biobank-scale genome-wide association studies (GWAS), recent epigenome-wide association studies (EWAS) have identified blood-derived DNA methylation associations within the *FKBP5* locus with neurodegenerative diseases^18^, as well as all-cause mortality^19^.

Preclinical studies have recently shown that FKBP51 is a crucial driver of persistent pain states following physical injuries and traumatic stress exposure^20–22^. Moreover, FKBP51 is up-regulated at spinal cord level in the early days after injury and spinal FKBP51 specifically maintains persistent pain of neuropathic and inflammatory origin, as its genetic deletion and spinal pharmacological inhibition provide significant pain relief^20, 23^. Crucially, we reported that early life trauma in mice leads to a reduction in spinal *Fkbp5* DNAm and prolonged subsequent pain states when exposed to inflammation in adulthood^23^. We therefore hypothesised that stress exposure promotes the vulnerability to chronic pain in an FKBP51-dependent mechanism. Here, we tested this hypothesis using a model of sub-chronic stress exposure in adult mice. Our results suggest that following stress exposure, FKBP51 drives the susceptibility to persistent pain in male mice at least. This increased vulnerability is accompanied by changes in DNAm in nociceptive signaling and stress-regulated genes at spinal cord level, including a reduction in DNAm in *Fkbp5.* However, complete reversal of these DNAm changes is not necessary to reverse the primed state.

## 2. Results

### 2.1 Sub-chronic stress primes for hyper-responsiveness to inflammatory pain

Male and female mice were exposed to the restraint stress paradigm (RS) 14 days before the injection of the inflammatory agent Complete Freund’s Adjuvant (CFA) (Fig.1A). Both male and female mice displayed mechanical hypersensitivity that peaked on the first day of testing (day 1 following last restraint = day 4) and were no different from control animals by day 10. (Fig.1B). There were no sex differences in the response to RS. On day 14, animals received an injection of CFA into the hindpaw. CFA induced a short-lasting hypersensitive state in control, non-stressed, mice (mechanical thresholds were no different from baseline thresholds by day 35 in male and female mice (one-way ANOVA: male controls, day 25 *vs* day 0: p=0.013, day 35 *vs* day 0: P>0.05; female controls, day 25 *vs* day 0: p=0.004, day 35 *vs* day 0: P>0.05)). However, stress exposure exacerbated the CFA-induced mechanical hypersensitivity in both male and female mice (RM-ANOVA: day 20 to day 80: RS mice vs Control mice: RS effect: F_1,12_=15.9, p=0.002; sex effect: F_1,12_=7.5, p=0.018; no RS effect x sex interactions). Mechanical thresholds were not only lower overall in both male and female mice exposed to RS (RS effects in males: day 20 to day 70: control vs RS: F_1,6_=8.6, p=0.026; in females: day 20 to day 80: control vs RS: F_1,6_=15.5, p=0.009), but the duration of the CFA-induced mechanical hypersensitivity was also extended. Indeed, mechanical thresholds returned to baseline thresholds by day 80 in RS male mice (male RS, last significant difference on day 70: day 70 vs day 0: p=0.03; day 80 vs day 0: P>0.05) and never returned to baseline for RS female mice (female RS, day 80 vs day 0: p=0.007). Stress exposure also had a greater impact on female mice as the thresholds of stressed female mice were lower than stressed male mice (RM-ANOVA: Females RS vs Males RS, day 20 to day 80: F_1,6_=9.2, p=0.023). In another subset of male mice, we assessed whether RS could exacerbate the hypersensitive state induced by another inflammatory agent and, indeed, RS exacerbated the response to interleukin 6 (IL6) injected in the hindpaw (Fig.S1).

**Figure 1:**
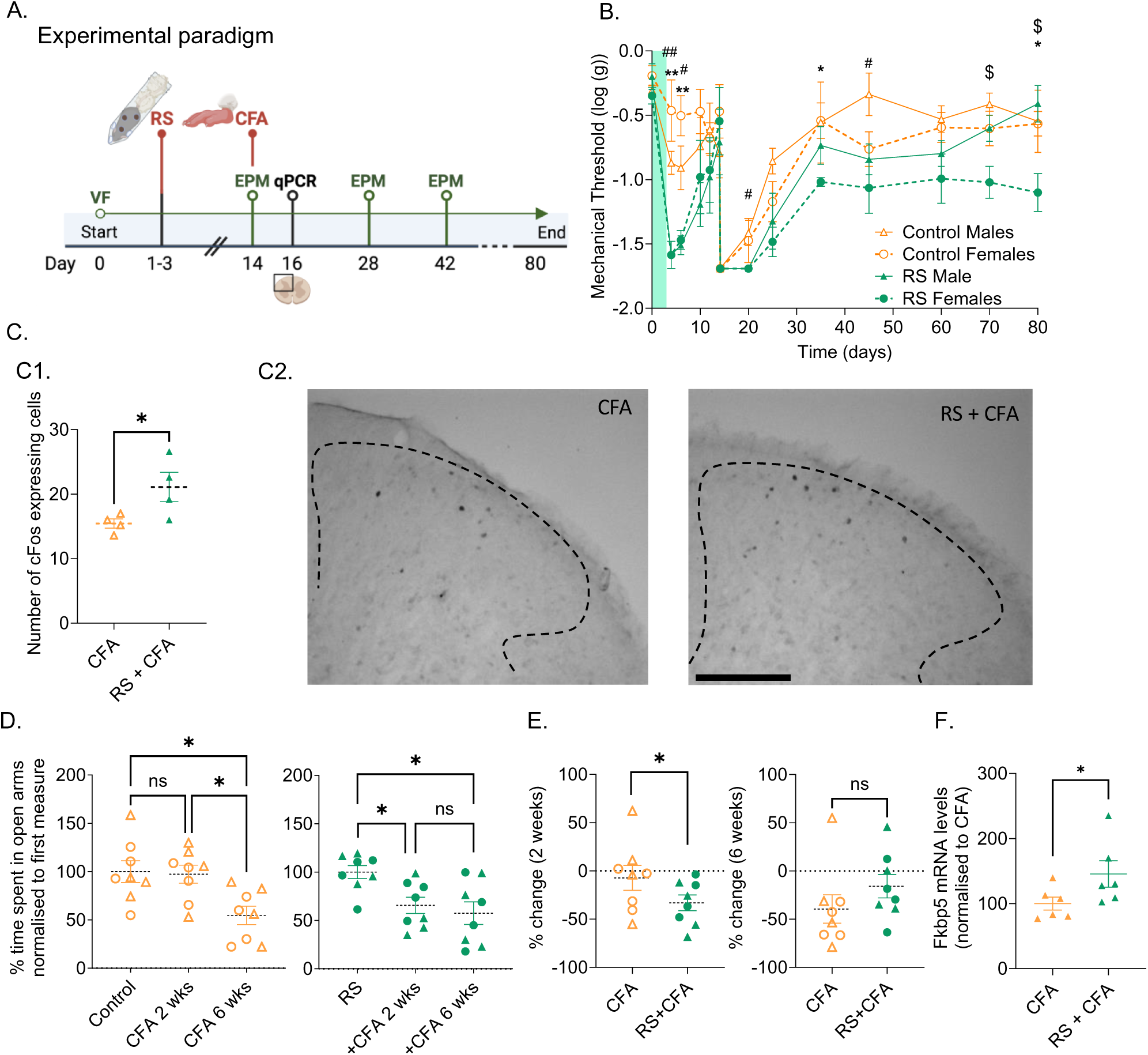
Sub-chronic stress primes for hyper-responsiveness to inflammatory pain. (**A**) Experimental design. (**B**) Hind paw mechanical withdrawal thresholds assessed in male and female mice using von Frey filaments. All mice received intra-plantar CFA injection on day 14. Post-hoc analysis: *: p<0.05: control females vs RS females; #: p<0.05 control males vs RS males; $ p<0.05: RS females vs RS males. Green panel indicates the 3 days of RS paradigm. (**C**) **C1**: number of c-Fos expressing cells in the ipsilateral superficial dorsal horn (lamina 1-2) 2 hours after CFA-injection; **C2**: Representative images of ipsilateral lumbar spinal c-Fos staining. Scale bar: 200µm. (**D**) Anxiety-like behaviour was measured using the elevated plus maze. Measures were taken at day 14, then at two- and six-weeks post-CFA (day 28 and day 44). (**E**) Anxiety-like behavior data from panel D, presented as a % change from baseline. There were significant differences in percentage change over time from pre- to two-weeks post CFA but not at two- to six-weeks post CFA, suggesting that RS accelerated CFA-induced anxiety-like behaviour. (**F**) RTqPCR quantification of *Fkbp5* mRNA levels. (**C-D**) *P<0.05.

While RS did not exacerbate the peak hypersensitivity after CFA, *i.e.* when it reached its maximum at 6h after CFA (day 14 plus 6h), it did promote the expression of cFOS measured 2h after CFA in the superficial laminae of the dorsal horn (Fig.1C), suggesting that stress exposure could promote nociceptive signalling. We next assessed the development of comorbid anxiety-like behaviors, often observed following stress exposure. Stress exposure accelerated the development of anxiety like behavior following CFA injection (Fig.1D, E). Animals who were not exposed to stress showed signs of anxiety only 6 weeks after CFA compared with stressed animals that showed signs at 2 weeks. Finally, we found that animals exposed to the RS paradigm had elevated levels of spinal *Fkbp5* mRNA compared to control animals 48h after CFA injection (day 16) (Fig.1F). This finding was similar to our previous observations of elevated spinal *Fkbp5* mRNA levels that occur in the first 48h of persistent injured state^21, 24^., suggesting that FKBP51 may be responsible for the longer lasting pain state observed in stressed mice.

### 2.2 FKPB51 drives stress-induced vulnerability to persistent pain in mice

We next tested the hypothesis that FKBP51 is an important contributor to the mechanisms driving the increased vulnerability to persistent pain observed following stress exposure. For this, we first used *Fkbp5* global knock out (KO) mice and exposed them to the RS plus CFA paradigm (Fig.1A).

When mice were exposed to the RS paradigm, we observed that the *Fkbp5* KO mice did not develop mechanical hypersensitivity as seen in WT mice. Both male and female KO mice had reduced RS-induced hypersensitivity (RM-ANOVA: day 0 to day 10: factor genotype: F_1,26_=15.9, p<0.001; no sex differences; post-hoc RM-ANOVA: day 0 to day 10: factor genotype: males only: F_1,13_=15.3, p=0.002; females only: F_1,13_=4.7, p=0.049). However, KO female mice were not different from WT female mice at any individual time point, suggesting that female KO were less resilient to RS than male KO. Following CFA injection, KO mice showed reduced mechanical hypersensitivity when compared to WT mice (RM-ANOVA, day 14 + 6h to day 60: factor genotype: F_1,26_=19.9, p<0.001; post-hoc males only: F_1,13_=22.0, p<0.001; post-hoc females only: F_1,13_=5.7, p=0.033). Unexpectedly, *Fkbp5* KO females were no different from WT females from day 45. Moreover, female KOs did not return to their baseline threshold for the duration of the observations (day 0 *vs* day 60: KO females threshold: p=0.024) while KO males had returned to their baseline thresholds by day 40 (day 0 vs day 40: KO males threshold: p>0.05). Overall, these results indicated that FKBP51 may be driving the stress-induced increased vulnerability to persistent pain in male mice at least. Therefore, we further explored the mechanisms of FKBP51-driven persistent pain vulnerability in male mice only.

### 2.3 FKBP51 inhibition during stress exposure prevents stress-induced increased vulnerability to persistent pain

To dissect the role of FKBP51 in promoting persistent pain during the stress phase and the injury phase of our paradigm, we used the specific FKBP51 inhibitor SAFit2. Our previous work had shown a delay in the initiation of FKBP51 antagonism when SAFit2 is encapsulated in a phospholipid gel for slow release (SAFit2-VPG). Under these conditions, SAFit2 is active for a maximum of 7 days^20, 21^. SAFit2-VPG was injected 2 days before the start of the RS paradigm to ensure complete FKBP51 blockade during the RS paradigm but not at the time of the CFA injection. While FKBP51 inhibition did not prevent the RS-induced mechanical hypersensitivity, it prevented the exacerbation to subsequent CFA-induced mechanical hypersensitivity (Fig.2B). Vehicle treated mice had returned to their baseline threshold by day 80 (last significant difference on day 70; day 0 *vs* day 70: p=0.012) whereas SAFit2-treated mice had returned to baseline threshold by day 35 (last significant difference on day 30; day 0 vs day 30: p=0.0032), following a time course similar to the WT mice non-exposed to RS (Fig.1B). Moreover, there was a significant difference in CFA-induced mechanical hypersensitivity between SAFit2-VPG and vehicle treated animals (Fig.2B; RM-ANOVA: factor treatment; day 15 to day 80: F_1,10_=16.7, p=0.002).

**Figure 2:**
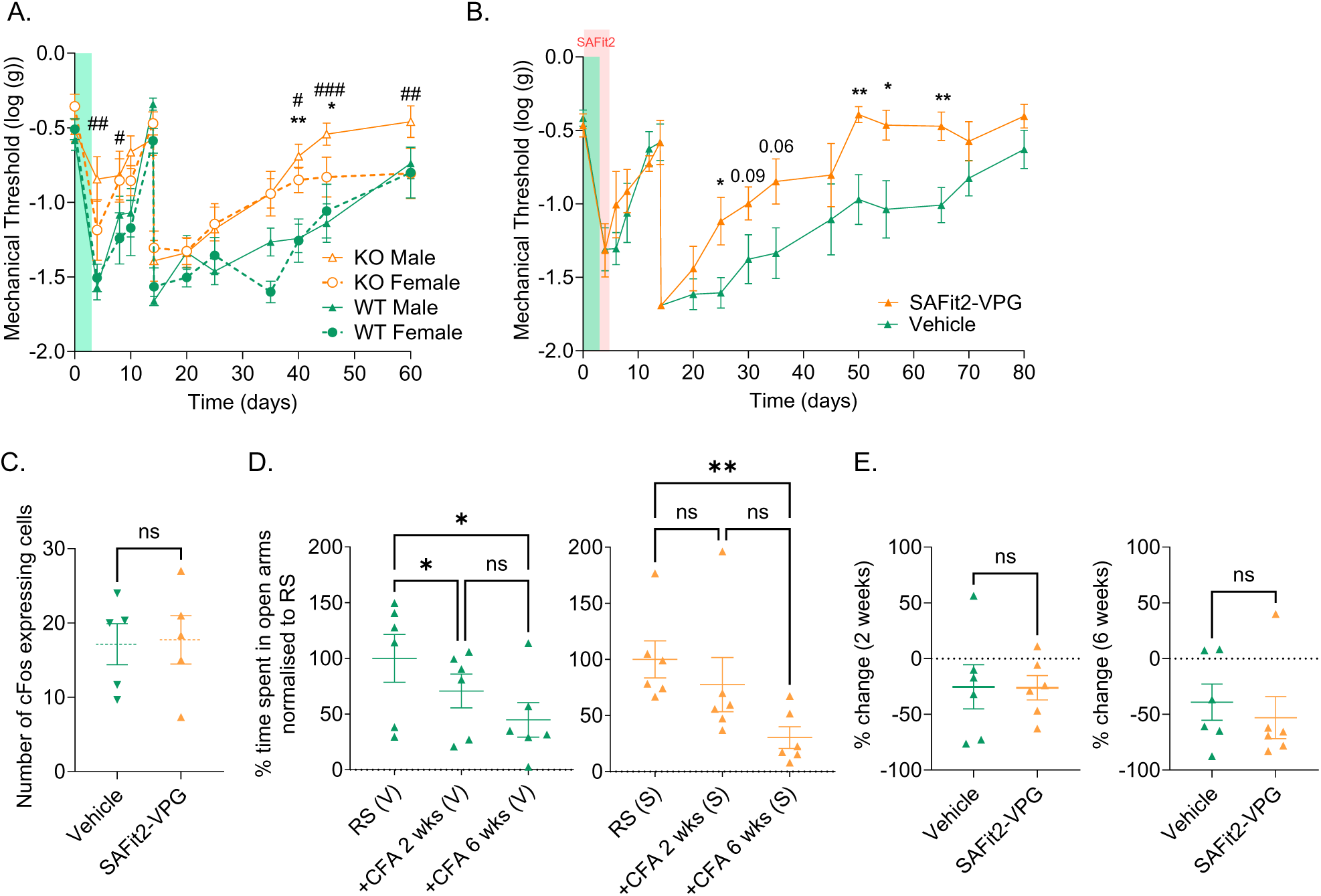
FKPB51 drives stress-induced vulnerability to persistent pain in mice. (**A**) Hind paw mechanical withdrawal thresholds measured using von Frey filaments in WT and KO *Fkbp5* mice. All mice received intra-plantar CFA on day 14. *Fkbp5* global deletion prevented the full development of RS hypersensitivity in male mice and to a lesser extent in female mice. Green panel indicates the 3 days of RS paradigm. Post-hoc analysis: *: WT females vs *Fkbp5* KO females; #: WT males vs *Fkbp5* males. (**B**) Hind paw mechanical withdrawal thresholds measured using von Frey filaments in male mice receiving either Vehicle- or SAFit2-VPG, administered two days prior to the onset of RS. Green panel indicates the 3 days of RS paradigm and red panel indicates the days of active SAFit2-VPG treatment. (**C**) Number of cFos expressing cells in the ipsilateral superficial dorsal horn (lamina 1-2) 2h after CFA injection. (**D, E**) Anxiety-like behaviour was measured using the elevated plus maze. (E) Anxiety-like behavior data from panel D, presented as a % change from baseline. **P<0.01, *P<0.05.

As we were surprised by the absence of effect of SAFit2 in the RS phase of the paradigm, we repeated this experiment with another SAFit2 formulation, injected i.p., twice a day for 5 consecutive days, to inhibit FKBP51 during the RS phase, as before. This delivery method increases SAFit2 plasma concentration at peak (personal communication). Again, we observed no effects of FKBP51 inhibition on the RS-induced mechanical hypersensitivity, while SAFit2 did reduce the subsequent CFA-induced hypersensitivity (Fig.S2A).

We next assessed the impact of FKBP51 inhibition during the RS phase on CFA-induced early nociceptive signaling. There was no difference between SAFit2-VPG and vehicle-VPG treated mice in cFos activation evoked by CFA (Fig.2C), implying that SAFit2 did not reduce the exacerbation of CFA-induced cFos by RS we had previously observed (Fig.1C).

Finally, we looked at the impact of FKBP51 inhibition during RS on anxiety-like behavior and found that SAFit2-VPG delayed the development of anxiety-like behavior, previously observed in wild type mice injected with CFA after stress exposure (Fig.1D, E and Fig.2D, E). Control experiments in naïve WT mice revealed that SAFit2-VPG alone did not modify anxiety-like behavior, measured using the EPM, or locomotion, as measured by the total distance travelled in the Open Field 5 day after SAFit2-VPG injection (Fig.S2B, 3C).

### 2.4 Sub-chronic stress induces rapid changes to stress signaling at spinal cord level

We next investigated whether stress exposure causes rapid changes to spinal cord signalling involved in the maintenance of long-term pain states. FKPB51 is upregulated by cortisol release and we found that in addition to mechanical hypersensitivity, the RS paradigm also led to an increase in blood serum glucocorticoids (Fig.3A, B), together with an upregulation of spinal *Fkbp5* mRNA measured 48h after the last restraint (day 5) (Fig.3C). There was also a downregulation of *Nr3c1*, the gene encoding for the glucocorticoid receptor (GR), particularly of the beta isoform which is associated with glucocorticoid resistance^25^ (Fig.3C). However, RS alone did not induce anxiety-like behaviour at the same time point (day 5 = 48h post RS; Fig.3D) and at day 14 (Fig.3E).

**Figure 3:**
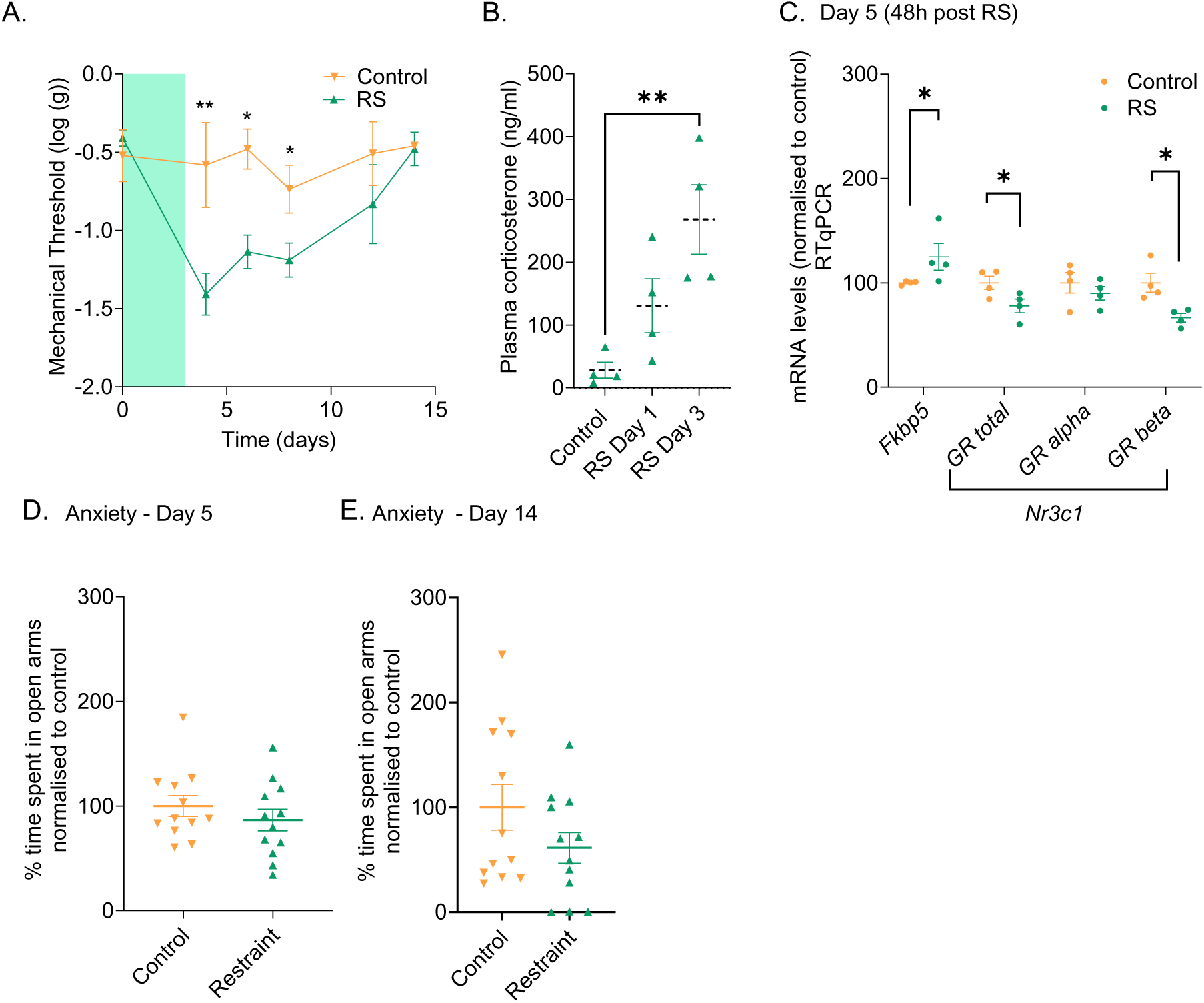
Sub-chronic stress induces rapid changes to stress signaling at spinal cord level. **(A)** Hind paw mechanical withdrawal thresholds assessed in male mice using Von Frey filaments. Green panel indicates the 3 days of RS paradigm. *p<0.05. Post-hoc one-way ANOVA. (**B**) Endogenous plasma CORT levels measured over the course of the RS paradigm. **P<0.01. (**C**) spinal mRNA levels assessed using RTqPCR at day 5. Graph shows percentage mRNA change normalized to control. (**D,E**) Anxiety-like behaviour assessed using EPM.

### 2.5 Sub-chronic stress induces long-lasting changes to the spinal methylome, including in the promoter of *Fkbp5*

We have previously shown that *Fkbp*5 DNAm is reduced in the rodent spinal cord both after injury^20^ and after early life adversity^23^, when reductions in DNAm may increase vulnerability to chronic pain later in life. We therefore assessed whether sub-chronic stress in adulthood modified the DNA methylome at spinal cord level and established to what extent the DNAm landscape of stress related genes such as *Fkbp5* may be modified. Spinal cord dorsal halves (*i.e.* ipsilateral + contralateral dorsal horns) were dissected on day 14 after the RS paradigm and prepared for both DNAm analysis and mRNA sequencing (RNAseq). At this time point, mice exposed to stress are no longer hypersensitive compared to control mice (Fig.4A), cortisol levels are no longer elevated (Fig.4B) but RS exposed mice are primed for hyper-responsiveness to CFA (Fig.1B). Differential DNA methylation analysis revealed that, of the 192,917 available post-QC CpGs, 11,644 probes were modulated by RS at a nominal significance of p < 0.05 (9,119 hypomethylated and 2,525 hypermethylated). However, no individual CpG surpassed an FDR threshold of p < 0.05 (Fig.S3A-B).

**Figure 4:**
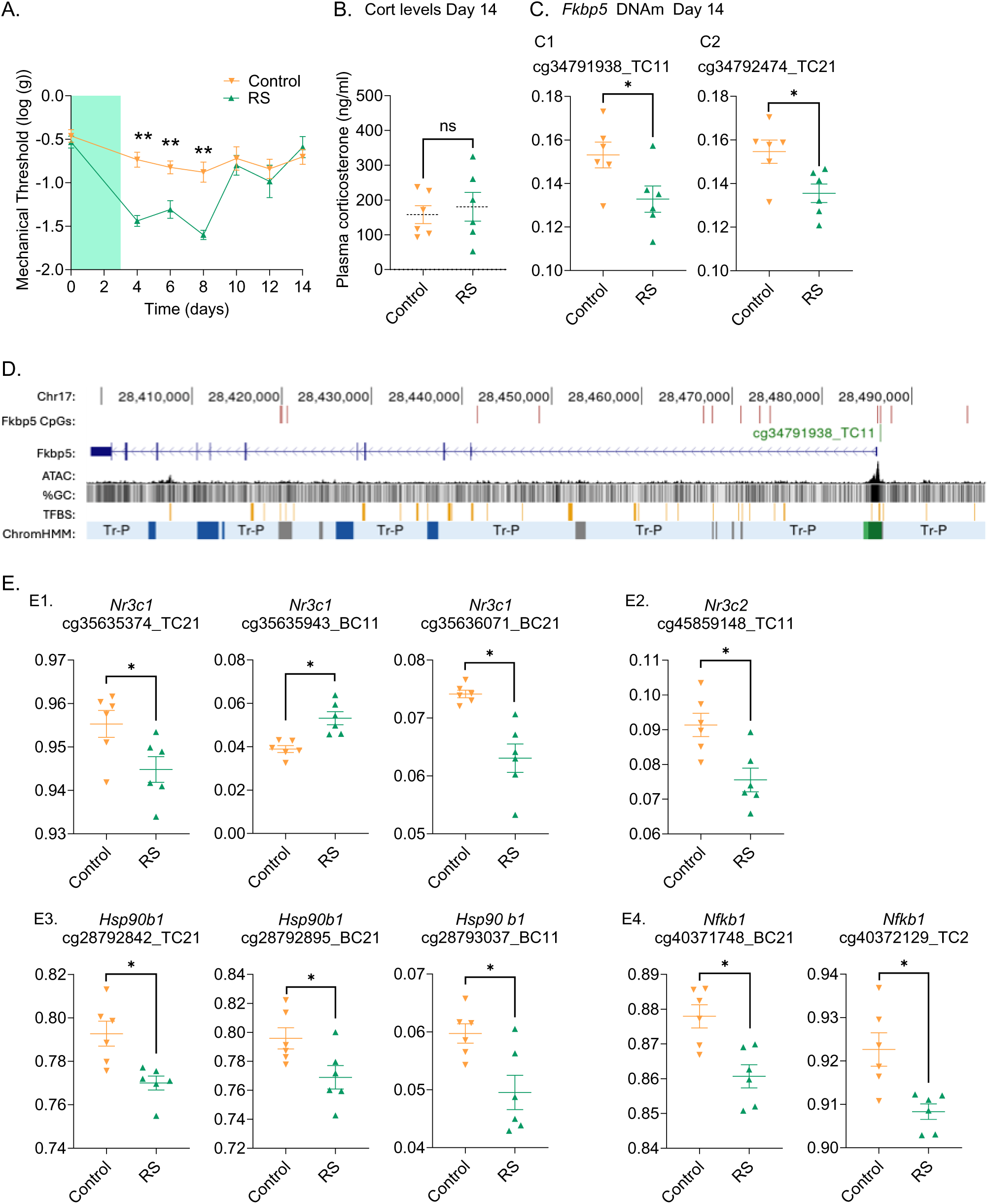
Sub-chronic stress induces long-lasting changes in the spinal DNA methylome, including in *Fkbp5* promoter region. (**A**) Hind paw mechanical withdrawal thresholds assessed in male mice using Von Frey filaments. RM ANOVA: T0 to T10: F_1,10_=45.4, p<0.001. Post-hoc analysis one-way ANOVA: **p<0.01 (**B**) Blood corticosterone levels from the same mice collected on day 14. (**C**) DNA methylation (DNAm) levels assessed at day 14, using the Infinium Mouse Methylation BeadChip arrays, at 2 independent CpG base. (**D**) Genomic region displaying the structure and regulatory elements associated with *Fkbp5*. The horizontal axis represents the genomic coordinates, spanning from 28.4 to 28.5mb on chromosome 17. The gene structure is shown with blue boxes indicating exons and arrows showing transcription direction. *Fkbp5* associated CpG sites present on the array beadchip are marked above the gene in red, highlighting potential methylation sites. A CpG (cg34791938_TC11), differentially methylated by RS, is indicated in green. Chromatin accessibility (ATAC sequencing), CG content, transcription factor binding sites and 15 state chromatin segmentation data derived from mouse neural tube tissue (E15.5) on the UCSC genome browser are displayed below the gene model. (**E**) DNA methylation levels in selected genes assessed using the Infinium Mouse Methylation BeadChip arrays. Y-axis: methylation levels (0-1). (**C,E**) * p<0.05 nominal significance.

We annotated the location and chromatin state according to mouse neural tube tissue^26^ of the 11,644 probes with nominal significance, to establish the importance that these probes may have on functional changes to gene expression. Hypomethylated CpGs in RS mice were enriched in active (Tss and TssFlnk) promoter sites (43.0% versus 24.1% array background, p <1.10^-4^) whereas these regions were underrepresented in CpGs hypermethylated by RS (21.0%, p = 4.10^-4^) (Fig.S3C). Consistent with these findings, Gene Set Enrichment via GREAT^27^ with the 11,644 probes satisfying nominal significance further confirmed an enrichment (49.19% versus 38.51% array background; p < 0.0001) of probes within 5 kb of the nearest TSS (Fig.S4D-E). Enrichments for biological processes were identified including ubiquitin homeostasis (>8 fold-enrichment, FDR p = 0.024), glutamate catabolism (>2.5 fold-enrichment, FDR p = 0.049) and retrograde axonal transport (>2 fold-enrichment, FDR p = 0.048). Additionally, heterotrimeric G-protein binding was identified as a molecular function (Table S1).

We next focused on stress related genes, including *Fkbp5*. A single probe (cg34791938_TC11) overlapping *Fkbp5* was revealed to be hypomethylated in RS relative to control mice (median log2FC = -0.24, 2% raw change in DNAm; control β: 0.15 ± 0.01 *vs* RS β: 0.13 ± 0.01; p = 0.037; Fig.4C1). This CpG is located within a CpG shore at chr17:28,486,463 in an active promoter region 313 bases upstream from the nearest TSS and resides in close proximity to several transcription factor binding sites (Fig.4D). There were no other significant changes in any of the other 13 CpGs located in the *Fkbp5* sequence (Table S2). However, a CpG (cg34792474_TC21) encoding a lincRNA (LOC102639076) associated with *Fkbp5* by GREAT and located proximal to several transcription factor binding sites (PITX2, TFAP2C, ZIC1, ZIC4 and ZIC5), 37 Kb upstream to the canonical TSS of *Fkbp5* (Chr17:28,486,150), was similarly hypomethylated in RS mice (median log2FC = -0.22; 2% raw change in DNAm; control β: 0.16 ± 0.01 vs RS β: 0.14 ± 0.01, p = 0.019; Fig.4C2). A number of HPA axis relevant genes also stood out as being differentially methylated by RS (at a nominal level), often in key regulatory regions, including *Nr3c1* (Fig.4E1), *Nr3c2* (Fig.4E2), *Hsp90b1* (Fig.4E3) and *Nfkb1* (Fig.4E4) (Table 1).

**Table 1:** Differentially Methylated Probes (DMPs) RS *vs* Control associated with genes linked to the HPA axis. CpGs are mapped to their closest gene, distance and chromatin state using mouse neural tube (E15.5) segmentation. FC: log2-fold change contrast. HMM: Chromatin state signatures established using hidden Markov model. Chr: Chromosome

| Probe ID | Location | Gene | chrHMM | CpG Island | Distance to TSS (b) | Log<sub>2</sub>FC | p |
| --- | --- | --- | --- | --- | --- | --- | --- |
| cg34791938_TC11 | chr17:28486463 | *Fkbp5* | Tss | S_Shore | -313 | -0.24 | 0.043 |
| cg31134886_TC11 | chr12:110696239 | *Hsp90aa1* | Tss | Island | 51 | -0.38 | 0.024 |
| cg34952220_BC21 | chr17:45572421 | *Hsp90ab1;Gm35399* | Tss | N_Shore | 835 | -0.22 | 0.015 |
| cg34952221_BC11 | chr17:45572453 | *Hsp90ab1;Gm35399* | Tss | N_Shore | 803 | -0.33 | 0.001 |
| cg31852209_BC21 | chr13:83530040 | *Mef2c* | EnhPois | OpenSea | 25800 | -0.35 | 0.009 |
| cg31852250_BC21 | chr13:83536465 | *Mef2c* | QuiesG | OpenSea | 32225 | -0.19 | 0.043 |
| cg31852551_TC11 | chr13:83575702 | *Mef2c* | QuiesG | OpenSea | 71462 | -0.19 | 0.043 |
| cg31852609_BC21 | chr13:83586607 | *Mef2c* | QuiesG | OpenSea | 82367 | -0.32 | 0.019 |
| cg31852653_TC21 | chr13:83592947 | *Mef2c* | QuiesG | OpenSea | 88707 | 0.29 | 0.016 |
| cg31853129_TC21 | chr13:83656519 | *Mef2c* | EnhPois | OpenSea | 152279 | -0.23 | 0.041 |
| cg31851968_TC21 | chr13:83504342 | *Mef2c;Gm33317* | Quies | OpenSea | 102 | -0.18 | 0.041 |
| cg40371748_BC21 | chr3:135637477 | *Nfkb1* | QuiesG | OpenSea | 31863 | -0.22 | 0.014 |
| cg40372129_TC21 | chr3:135667764 | *Nfkb1* | QuiesG | OpenSea | 1576 | -0.28 | 0.012 |
| cg35635374_TC21 | chr18:39422515 | *Nr3c1* | TxWk | OpenSea | 67332 | -0.32 | 0.024 |
| cg35635943_BC11 | chr18:39490039 | *Nr3c1* | Tss | Island | -192 | 0.49 | 0.001 |
| cg35636071_BC21 | chr18:39491544 | *Nr3c1* | Tss | Island | -869 | -0.26 | 0.007 |
| cg45859148_TC11 | chr8:76900117 | *Gm10649; Nr3c2* | Tss | Island | 57 | -0.31 | 0.008 |
| cg41489811_TC21 | chr4:132131095 | *Oprd1* | QuiesG | OpenSea | 13392 | -0.21 | 0.021 |
| cg28138698_BC21 | chr10:6979665 | *Oprm1;lpcef1* | TssFlnk | OpenSea | 801 | -0.27 | 0.005 |
| cg30318030_BC21 | chr12:3959937 | *Pomc* | Tss | Island | 4986 | -0.21 | 0.012 |
| cg39323159_BC21 | chr2:169633030 | *Tshz2* | Tss | N_Shore | -646 | -0.45 | 0.002 |
| cg39323590_TC21 | chr2:169662164 | *Tshz2* | QuiesG | OpenSea | 28488 | -0.22 | 0.031 |
| cg39325299_TC21 | chr2:169783649 | *Tshz2* | QuiesG | OpenSea | 149973 | -0.22 | 0.029 |
| cg39326195_TC21 | chr2:169842640 | *Tshz2* | EnhPois | OpenSea | 208964 | -0.21 | 0.014 |
| cg39326879_TC11 | chr2:169884840 | *Tshz2* | Tx | OpenSea | 251164 | 0.27 | 0.042 |
| cg39326881_BC21 | chr2:169884910 | *Tshz2* | Tx | OpenSea | 251234 | -0.24 | 0.032 |
| cg39326933_TC11 | chr2:169886400 | *Tshz2* | TxWk | OpenSea | 252724 | 0.23 | 0.017 |
| cg39327361_BC21 | chr2:169912922 | *Tshz2* | EnhPois | OpenSea | 279909 | -0.36 | 0.005 |

### 2.6 Sub-chronic stress induced-changes in the spinal DNA methylome are not fully reversed by FKBP51 inhibition despite its behavioural impact

As FKBP51 inhibition with the inhibitor SAFit2 prevented the exacerbation of CFA-induced mechanical hypersensitivity, we asked whether SAFit2 could prevent the RS-induced changes in spinal DNAm, including at *Fkbp5* cg34791938_TC11. As previously observed, mice treated with SAFit2-VPG during the RS paradigm developed mechanical hypersensitivity to the same extent as vehicle treated mice (Fig.5A). There was no difference in *Fkbp5* mRNA levels between vehicle and SAFit2-VPG treated mice at day 5 (48h post RS) (Fig.5B) suggesting that SAFit2-VPG did not prevent the upregulation of *Fkbp5* mRNA induced by the RS paradigm we had previously observed (Fig.3C).

**Figure 5:**
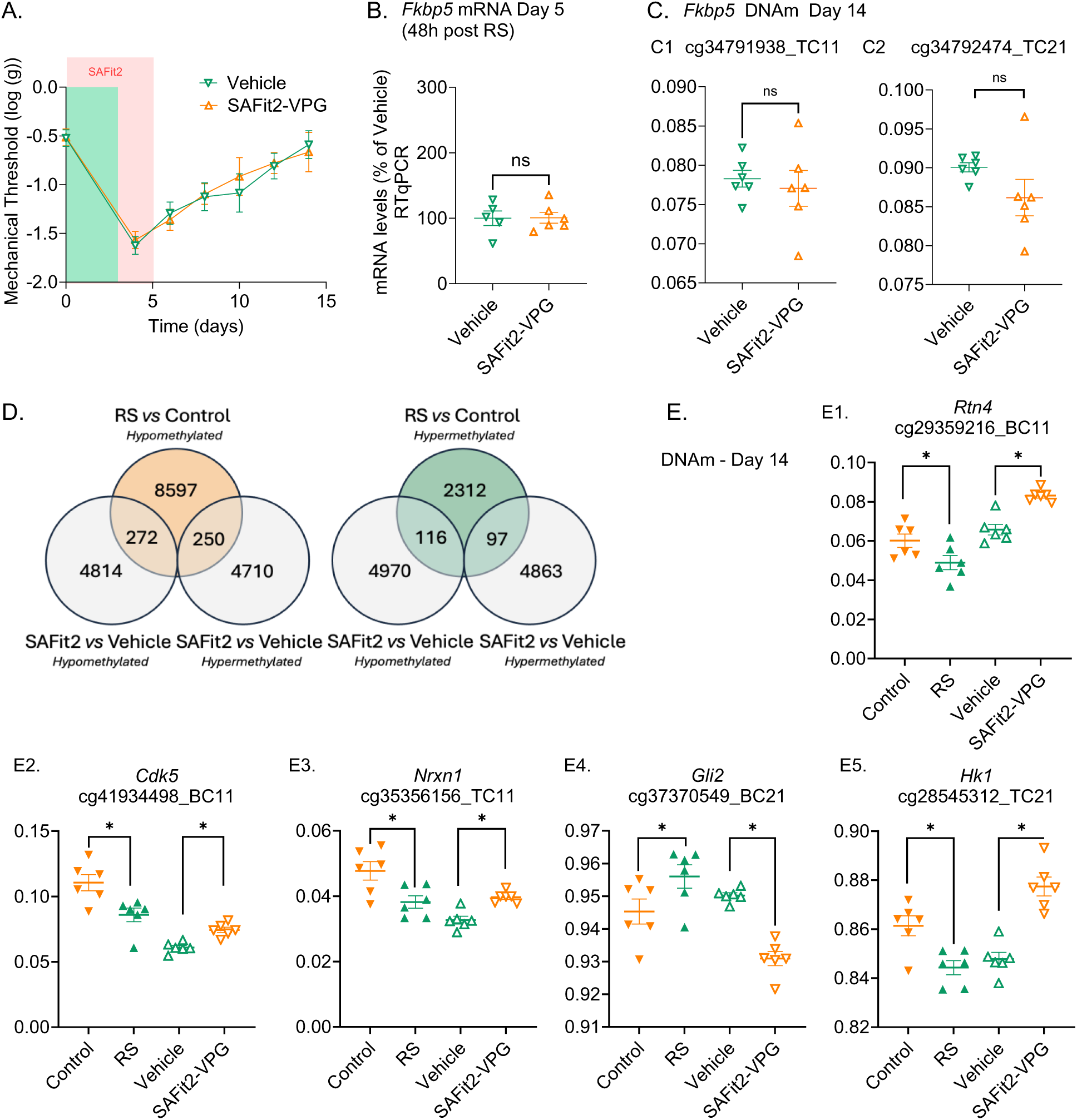
FKBP51 inhibition during stress exposure reverses some, but not all, changes in sub-chronic stress induced changes in spinal DNA methylome. (**A**) Hind paw mechanical withdrawal thresholds assessed in male mice using von Frey filaments. Green panel indicates the 3 days of RS paradigm and red panel indicates the days of active SAFit2 treatment. (**B**) *Fkbp5* mRNA levels in spinal cord superficial dorsal horn assessed by RTqPCR at day 5 (48h post RS). (**C**) *Fkbp5* DNAm status assessed using the Infinium Mouse Methylation BeadChip array. (**D**) Venn Diagram showing the intersection of hypomethylation and hypermethylated CpGs across the 2 DNAm studies. (**E**) DNA methylation levels in selected genes assessed using the Infinium Mouse Methylation BeadChip arrays. Y-axis: methylation levels (0-1). *p<0.05 nominal significance.

Using the same approach as in our first DNA methylation study, we found that 4,960 CpGs out of 191,134 total analytically robust CpGs were hypermethylated in SAFit2-VPG treated mice, whereas 5,086 CpGs were hypomethylated at a nominal p value threshold (p< 0.05; total 10,046 DMPs). There were similarly no individual CpGs surpassing a Benjamini-Hochberg FDR threshold of p < 0.05 (Fig.S3F-G). Probe location and chromatin segmentation in mouse neural tube tissue of the 10,046 CpGs did not significantly differ from the array background (Fig.S3H), nor were there any statistically significant terms identified by Gene Set enrichment via GREAT. Moreover, unlike the contrast between RS and control mice, which revealed enrichment of probes within 5 kb of the TSS (Fig.S3D-E), there was not an enrichment in probes within 5 kb of TSS for the 10,046 CpGs differing between RS mice treated with SAFit2 or vehicle (Fig.S4I-J).

There was also no difference in DNA methylation at *Fkbp5* cg34791938_TC11 (median log2FC = -0.031; <0.01% raw change in DNAm; vehicle β: 0.078 ± 0.001 vs SAFit2 β: 0.077 ± 0.006, p = 0.64; Fig.5C1) or cg34792474_TC21 (median log2FC = -0.077; <1% raw change in DNAm; vehicle β: 0.090 ± 0.001 vs SAFit2 β: 0.086 ± 0.006, p = 0.14; Fig.5C2) between vehicle and SAFit2-VPG treated mice exposed to RS. Taken together, these observations suggest that inhibition of FKBP51 during the RS phase may not revert changes to the *Fkbp5* DNAm landscape induced by the RS paradigm.

We also looked more widely at the intersection between the DMPs identified in the two DNAm studies to identify any DMPs regulated in opposite direction, that may contribute to the diverging pain phenotypes observed following RS with or without SAFit2 exposure. Intersections between hypermethylated and hypomethylated CpGs from each contrast revealed 366 CpGs regulated in opposite direction (250 CpGs hypomethylated RS *vs* Control and hypermethylated SAFit2 *vs* Vehicle and 116 hypermethylated RS *vs* Control and hypomethylated SAFit2 *vs* Vehicle, Fig.5D). These associated with 284 distinct genes and many of these (108/284) were oppositely regulated in active promoter regions (Tss and TssFlnk). Of particular interest were genes related to nociceptive signalling (Table 2), including *Rtn4* (Fig.5E1), which encode the protein Nogo-A, a neurite outgrowth inhibitor, whose inhibition has been proposed a novel approach for pain management^28^; *Cdk5* (Fig.5E2), that encodes for the cyclin-dependent kinase 5, also considered a potential target for the management of chronic pain^29^; *Nrxn1* (Fig.5E3), which encodes the neurexin 1 protein, previously linked to hypersensitive states^30^ and *Gli2* (Fig.5E4), previously linked to neuropathic pain^31^. We also observed interesting DNAm reversal in regions linked to quiescent state, *e.g.* in *Hk1* (Fig.5E5), that encodes the hemokinin 1, which binds to the NK1 receptor and is an important pain mediator^32^.

**Table 2:**
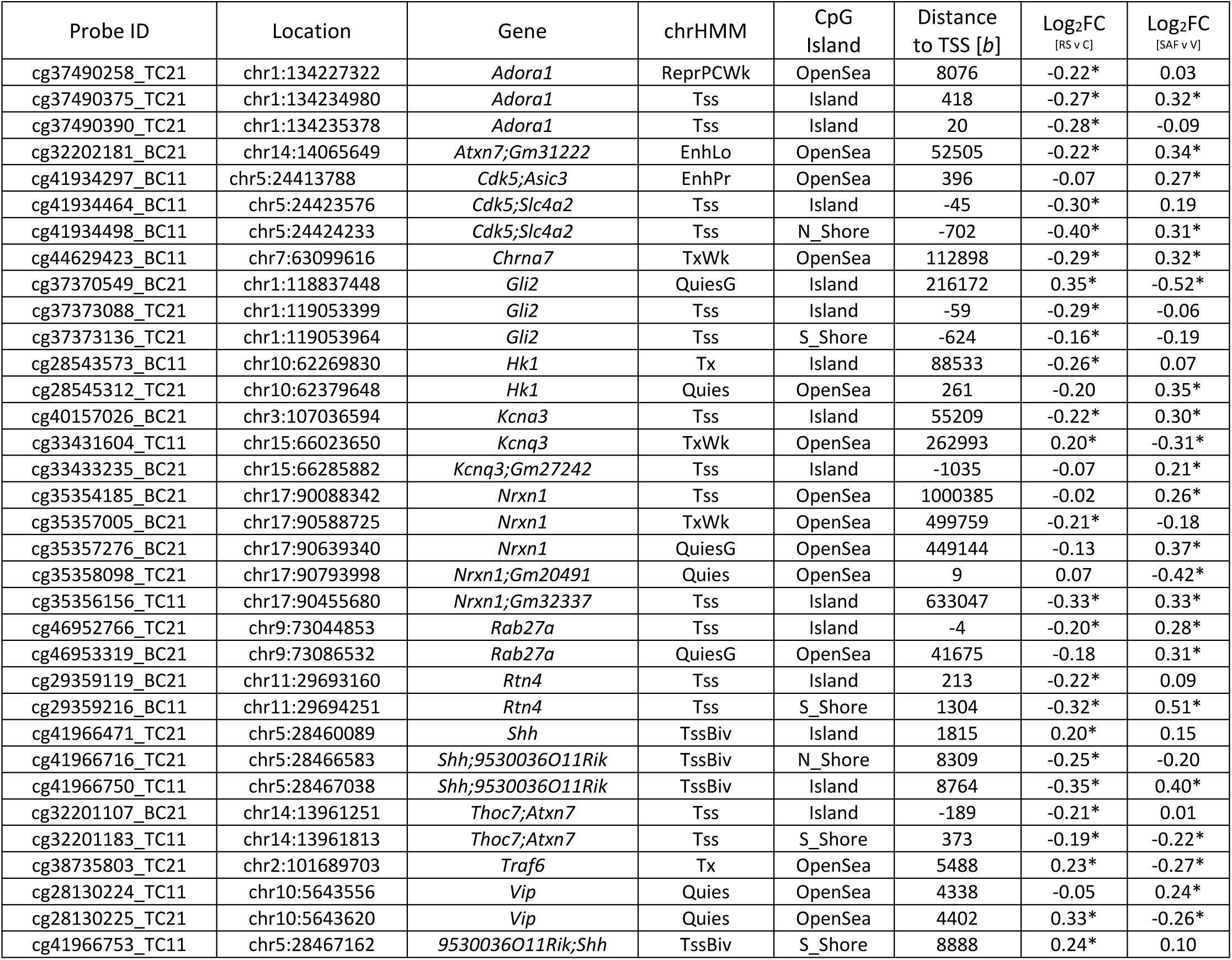
Inversely Differentially Methylated Probes (DMPs) in RS *vs* Control and SAFit2 *vs* Vehicle studies, associated with genes relevant to priming mechanisms and nociceptive processing. CpGs are mapped to their closest gene, distance and chromatin state using mouse neural tube (E15.5) segmentation. FC: log2-fold change contrast. HMM: Chromatin state signatures established using hidden Markov model. Chr: Chromosome. *: p<0.05 nominal significance.

### 2.7 Following restraint stress (RS) the *Fkbp5* loci CpGs cg34791938_TC11 and cg34792474_TC21 are hypomethylated and *Fkbp5* mRNA is upregulated

Differential gene expression (RS – Control) on log2-transformed RNA count intensities (n = 15533) revealed a difference in expression for 230 genes (153 downregulated genes and 77 upregulated genes) exceeding the criteria of FC > 1.2 (nominal p < 0.05). These genes were not significantly overrepresented in any biological process ontologies at an FDR corrected p value < 0.05. However, interestingly, *Fkbp5* was among the genes upregulated (1.3-fold of control) by RS (Fig.6A-C). These observations suggested that reduced DNAm after RS at both cg34791938_TC11 and cg34792474_TC21 could have some functional relevance to *Fkbp5* expression.

**Figure 6:**
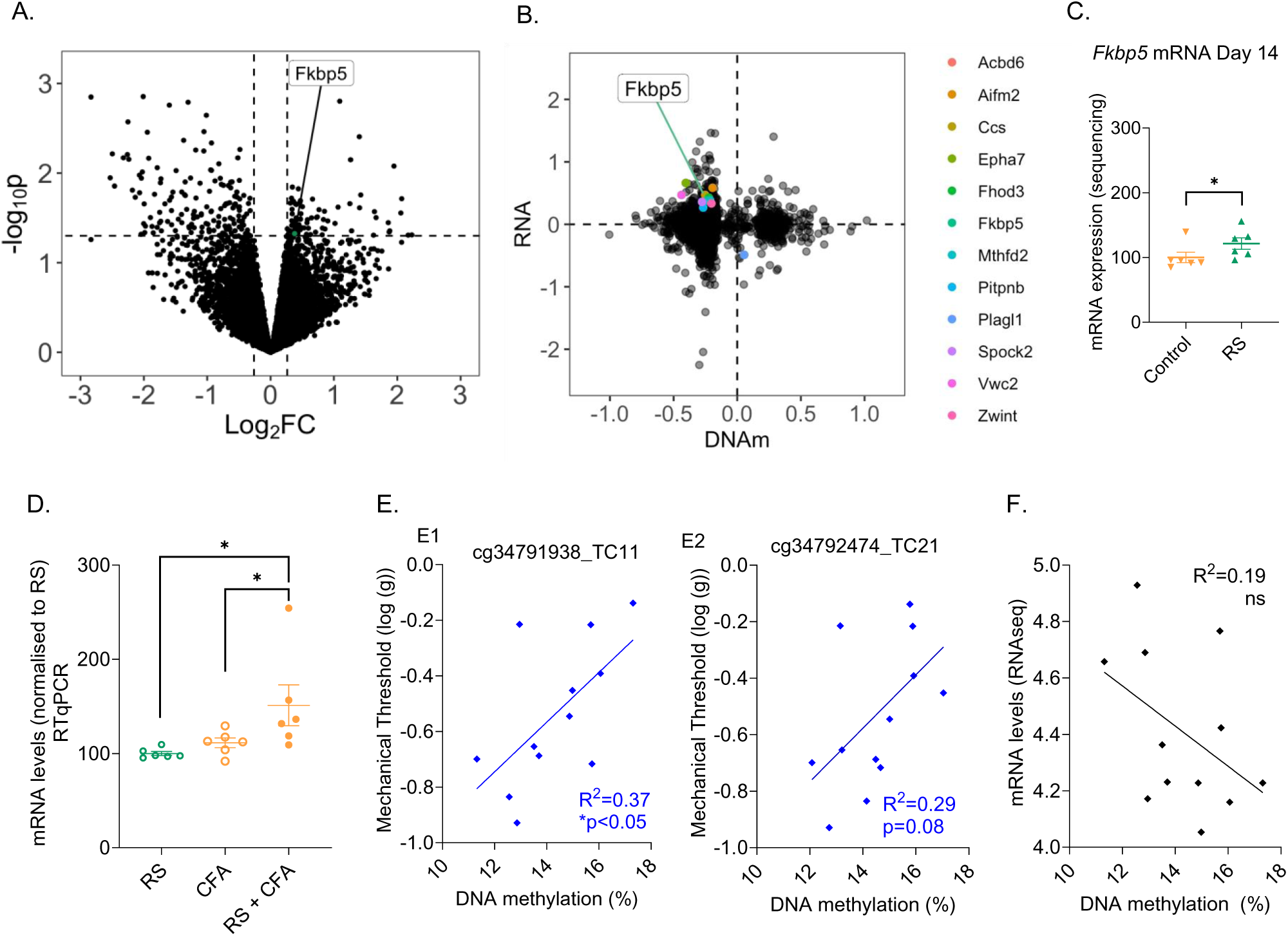
*Fkbp5* mRNA is upregulated after RS when both cg34791938_TC11 and cg34792474_TC21 are hypomethylated. (**A**) Volcano plot visualising the differential expression contrasting RS mice *versus* controls. The x-axis denotes the log2 fold change (FC) in RNA expression between the two conditions, with positive values indicating higher expression in the RS group and negative values indicating higher expression in controls. The y-axis represents the -log10 transformed p-values. Data are adjusted for cell type heterogeneity. (**B**) Scatter plot visualising the intersection between CpGs occurring in promoter regions that are differentially methylated by RS and corresponding changes to RNA expression in the same samples. Y-axis: Log_2_RNA expression (RS/control). (**C**) *Fkbp5* mRNA levels from sequencing data. (**D**) RTqPCR quantification of *Fkbp5* mRNA in the ipsilateral dorsal horn of the spinal cord 48h after CFA injection. *p<0.05. (**E**) Correlation between mechanical threshold (average day 4 to 10) and DNAm at cg34791938_TC11 (E1) and cg34792474_TC21 (E2). (**F**) Correlation between DNAm at cg34791938_TC11 and *Fkbp5* mRNA levels at day 14.

To investigate whether this reduction in DNAm could also be mechanistically associated with the elevated levels of *Fkbp5* mRNA seen in RS exposed animals 48h after CFA compared with non-stressed controls (Fig.1F), we compared spinal *Fkbp5* mRNA levels between RS alone, CFA alone and RS + CFA exposed animals and found that RS potentiated the expression of *Fkbp5* after CFA (Fig.6D), supporting the idea that RS exposure had primed the gene for responsiveness. We additionally found that the DNAm levels at cg34791938_TC11 correlated with the change in mechanical thresholds observed following the RS paradigm (cg34791938_TC11: r2 = 0.37, p < 0.05; cg34792474_TC21: r2=0.29, p=0.08, Fig.6E1, E2) but not with *Fkbp5* mRNA levels at day 14 (Fig.6F). There was no correlation between the *Fkbp5* mRNA levels at day 14 and the RS-induced change in mechanical thresholds, suggesting that long-term *Fkbp5* mRNA levels after the stress exposure were not driven by the response to RS (Fig.S4).

### 2.8 Exposure to stress in adulthood leads to epigenetic changes at spinal cord levels often not associated with changes in gene expression

The DNAm analysis for our study 1 (RS vs Control) indicated that 11,644 CpGs differed in RS mice compared with control and many (37.99%) occurred in actively transcribed promoter regions (Tss, TssFlnk) within their closest gene in mouse neural tube chromatin segmentation^26^. This included 509 of the 2,525 hypermethylated CpGs and 3,915 of the 9119 hypomethylated CpGs. When the CpGs occurring in promoters were paired against their corresponding RNA expression (Fig.6B), we found that 307 unique genes were associated with hypermethylated promoter CpGs and 2780 unique genes had hypomethylated promoter CpGs on day 14 after the RS procedure, when animal behaviour had returned to normal. However, only 1 of the hypermethylated genes (*Plagl1*) was concurrently down-regulated at day 14, and 11 of the hypomethylated genes (*Acbd6*, *Aifm2*, *Ccs*, *Epha7*, *Fhod3*, *Fkbp5*, *Mthfd2*, *Pitpnb*, *Spock2*, *Vwc2* and *Zwint*) were up-regulated (out of a total of 77 up-regulated genes). Taken together, these results suggested that the RS-induced changes in the epigenomic landscape were more likely to modify gene responsiveness to future challenges, rather than maintaining genes in a persistent up-regulated or down-regulated state.

## 3 Discussion

This study investigated the impact of stress exposure on subsequent inflammatory pain sensitivity using a sub-chronic stress paradigm (1h restraint on 3 consecutive days, RS) followed by injection of the inflammatory agent Complete Freund’s Adjuvant (CFA) into the hindpaw. The results demonstrate that exposure to stress primes both male and female mice for heightened and prolonged inflammatory pain, with a significant role played by the FKBP51 protein in mediating this stress-induced vulnerability to persistent pain, at least in male mice. Stress exposure induced numerous changes in DNAm at spinal cord level, including hypomethylation at a CpG overlapping the TSS of the gene *Fkbp5*, but reversal of hypomethylation at this locus was not necessary to prevent the increased vulnerability to persistent pain. Nonetheless, pharmacological inhibition of FKBP51 prevented the priming and reversed a number of DNAm changes identified in regulatory sequences of stress and nociceptive signalling related genes, suggesting that FKBP51 and the activation of the HPA axis was key to the underlying mechanisms.

### Sex differences in stress-induced increased vulnerability to persistent pain

Male and female mice exposed to sub-chronic stress showed mechanical hypersensitivity following stress and exacerbated hypersensitivity following CFA injection compared to non-stressed mice. However, the stress-induced increase in CFA-induced hypersensitivity was greater and longer-lasting in females, indicating a sex-dependent exacerbation of pain following stress. This aligns with existing literature suggesting that females often exhibit greater sensitivity to chronic pain and stress-related disorders^33–35^. Moreover, *Fkpb5* KO females were less resilient than males to both stress-induced hypersensitivity and stress-induced increased persistent pain vulnerability. This was surprising as FKBP51 is known to drive stress persistent pain equally in both male and female mice^20–22^. Overall, our initial findings suggested that FKBP51 was likely to drive the increased susceptibility to persistent pain in male mice at least. As our statistical analysis did not indicate significant sex x treatment interactions, our study may have been under-powered to identify significant changes in female mice.

### FKBP51 as a mediator of stress-induced persistent pain vulnerability but not sub-chronic stress induced hypersensitivity

To confirm the role of FKBP51 in the increased susceptibility to persistent pain, we used the specific inhibitor SAFit2, as before^20, 21^. While both male and female *Fkbp5* KO mice presented with reduced stress-induced hypersensitivity, pharmacological inhibition of FKBP51 with SAFit2 did not reduce the hypersensitivity induced by this sub-chronic stress exposure. These results were surprising, as SAFit2 can prevent the mechanical hypersensitivity following acute prolonged stress exposure in a rodent model of Post-Traumatic Stress Disorder^22^. In this paradigm, the stress exposure lasts 2h and does not re-occur on multiple days as in our paradigm. Our data therefore suggest that inhibition of FKBP51 alone cannot prevent the decrease in threshold induced by sub-chronic stress. Nonetheless, FKBP51 inhibition was sufficient to prevent the stress-induced exacerbation of the subsequent CFA-induced pain state.

### FKBP51 and Stress Signaling

Stress exposure led to an increase in blood serum glucocorticoids. This was accompanied by an increase in spinal *Fkbp5* mRNA and a decrease in spinal *Nr3c1* mRNA, the gene encoding the glucocorticoid receptor (GR), 48h after the last exposure to stress. While changes in expression of stress-activated genes occur within minutes of stress exposure following the orchestrated release of numerous stress mediators^36–38^, we chose a later time point for our analysis of gene expression. This is because we were interested in the link with the persistent pain phenotype and had previously reported that *Fkbp5* mRNA was elevated for at least up to 48h after the initiation of persistent pain states^20, 24^. We had found that *Nr3c1* mRNA was upregulated at this time point, while here we report a downregulation following stress exposure, suggesting differences in the regulation of the HPA axis between sub-chronic stress exposure and physical injury. The observations from the current study fit with previous reports of reduced GR expression in humans following NR3C1 increased DNAm in the promoter region upon substantial stress exposure^39–43^. We also report an increase in *Nr3c1* DNAm at cg35635943_BC11 within 200 bp of TSS (-192 bases) following stress exposure, suggesting that this was likely to drive the early downregulation in *Nr3c1* gene expression observed at day 5. *Nr3c1* mRNA was no longer downregulated at the time point of the DNAm analysis, suggesting that the stress-induced change in epigenome was longer lasting than the change in gene expression. Crucially, the FKBP51 inhibition that prevented the primed state did not reverse the stress-induced change in *Nr3c1* or in *Fkbp5* DNAm.

Stress exposure also led to an increase in CFA-induced cFos in Laminae I-II of the superficial dorsal horn, indicating enhanced nociceptive signalling. While FKBP51 inhibition prevented the RS-induced priming, it did not have any effect on cFos expression, suggesting that prevention of early cFos expression is not required to prevent the prolongation of persistent pain states. This aligns with our previous findings, showing that *Fkbp5* KO mice have reduced mechanical hypersensitivity in persistent pain states but similar levels of spinal cFos expression to wild-type mice 2h after pain state induction^20^. These observations also confirmed the lack of correlation between cFos expression and pain behaviour reported by others^44^. It also suggests that *Fkbp5* deletion and pharmacological inhibition does not reduce primary afferent input into the superficial dorsal horn following noxious stimulation, which confirms our hypothesis that FKBP51 drives persistent pain at the level of the central nervous system^20^.

### *Fkbp5* DNAm and chronic pain vulnerability

We and others have shown that early life trauma can modify *Fkbp5* DNA methylation landscape. In particular, we reported that early life trauma could reduce *Fkbp5* DNA methylation at spinal cord level, where FKBP51 has been shown to drive persistent pain states. Here, we report that exposure to sub-chronic stress in adulthood also leads to a reduction in DNAm in *Fkbp5* (cg34791938_TC11; Chr17:28486463, -313 bases from TSS) at spinal cord level. This is an important observation, as it is currently unclear whether stress may have the same impact in young people and adults, with studies supporting the idea that adults may be more resilient to intense stress exposure^17^. It was also crucial for our hypothesis to demonstrate that stress exposure in adulthood can lead to DNAm changes at spinal cord level, a tissue rarely investigated in human studies.

The nominal change in DNA methylation we report seems to prime *Fkbp5* for hyper-responsiveness to subsequent challenges, as stressed mice had a higher level of *Fkbp5* mRNA after CFA injection when compared with mice receiving only CFA and mice that were only stressed. Together, these observations would suggest that exposure to stress primes for chronic pain vulnerability through de-methylation of *Fkbp5*. However, findings from this study also suggest that de-methylation of *Fkbp5* alone is not sufficient to promote vulnerability, as inhibition of FKBP51 during the stress exposure period prevented the priming but not the change in DNAm in *Fkbp5*. Nonetheless, FKBP51 inhibition with SAFit2 did reverse a number of stress-induced changes DNAm (Table 2), notably in the pro-nociceptive genes *Rtn4, Cdk5, Nrxn1* and *Gli2*.

### Epigenetic changes and long-term pain vulnerability

Our experiments suggest that stress exposure leads to more long-lasting changes to DNAm than mRNA expression levels. Indeed, we report 4,424 DMPs within promoter active chromatin overlapping 3,590 unique genes (3,087 of these available in corresponding RNA sequencing data), 14 days after stress exposure. At this time point, differences in gene expression between stress exposed and control mice were very few (n=230, with Fold-change > 1.2 and nominal p value < 0.05). These results suggested that the expression of a significant number of genes was not impacted by the changes to the DNAm landscape alone, even when potential cell heterogeneity effects were considered, and that they may have been primed by the sub-chronic stress exposure. This aligns with the observations that DNA methylation, chromatin dynamics and transcription factor occupancy work on differing time scales^45^ and that DNA methylation may contribute to the stability of transcriptional regulation^46^.

On the other hand, our findings support the idea that DNA methylation profiling may offer valuable insights into various biological processes and have the potential to reveal clinically relevant information, as methylation profiles may reflect not only the activity of transcription factors but also serve as potential biomarkers for disease states. Recent studies have demonstrated that even non-affected tissue can yield informative DNA methylation profiles, aiding in treatment decisions, clinical cohort stratification, and potentially guiding personalized medicine^47^. Large-scale cohort studies promise to deepen our understanding of how specific DNA methylation patterns relate to disease phenotypes, enhancing the potential for methylation to be used in disease prognosis^46^.

### Conclusion: implications for chronic pain and stress disorders

Our findings indicate that sub-chronic stress primes both male and female mice for prolonged inflammatory pain, with FKBP51 proving crucial to this vulnerability, particularly in male mice. The molecular insights gained, including DNAm changes in pain-related regulatory gene sequences at spinal cord level, underscore FKBP51’s potential as a therapeutic target for chronic pain in humans.

## 4 Material and Methods

Extended methods can be found in the supplementary file.

### 4.1 Animals

ARRIVE guidelines were followed. Wild-type (WT) experiments were performed using 8–10-week-old C57BL/6 mice of both sexes (N=146; Charles River, UK). Male and female *Fkbp5 +/-* mice (N=16) and their WT littermates (N=14) were bred in house and genotyped according to Maiaru *et al.*^20^. Mice were housed in individually ventilated home cages in a temperature-controlled environment with a 12-hour light-dark cycle, with food and water available ad libitum. All procedures were carried out in accordance with the United Kingdom Animal Scientific Procedures Act 1986.

### 4.2 Sub Chronic Stress (restraint stress)

Mice were restrained for 1 hour/day for three consecutive days in 50ml falcon tubes, adapted to allow ventilation. Restraint onset was standardised to 10am to control natural fluctuations in circulating CORT^48^ that occur through the day^49^. Where possible, restrained mice were separated from controls to prevent social transmission of stress through odour^50^.

### 4.3 Models of inflammation

Acute hind paw inflammation was induced by injection of the pro-inflammatory cytokine interleukin 6 (IL-6; Sigma), while a longer lasting inflammatory pain state was induced by intra-plantar administration of Complete Freund’s Adjuvant (CFA; Sigma). Briefly, either 25µl of 0.1ng IL6 or undiluted CFA, was injected subcutaneously into the plantar surface of the left hind paw, under light anaesthetic (2% isoflurane in oxygen), in line with previous work from the lab^20^.

### 4.4 Drugs

The FKBP51 inhibitor, SAFit2, was synthesised as previously described^51^. The drug was used at a concentration of 200mg/kg, dissolved in two vehicles (see supplementary for vehicle and delivery methods)^21^. SAFit2-VPG was administered two days prior to the first restraint. According to previous experiments in the lab, the gel formulation can provide slow drug release lasting up to 7 days, thus SAFit2-VPG administration supplied FKBP51 inhibition for the duration of the restraint^21, 52^. Control mice received injection of the vehicle formulation only. Drug group was blinded by a member of the lab.

### 4.5 Mechanical withdrawal thresholds

Hind paw mechanical withdrawal thresholds were assessed using Von Frey fibres applied to the plantar region of the left hind paw via the up-down method^53, 54^., at a starting force of 0.6g (Ugo Basile SRL). Mice were habituated for 45 minutes before testing and measures were completed at the same time each day. Thresholds have been log-transformed in accordance with Mills *et al.* ^55^, to account for the logarithmic distribution of fibres.

### 4.6 Elevated plus Maze (EPM)

The elevated plus maze^56–58^ was used to measure anxiety-like behaviour. Mice were placed on the centre of the EPM (Ugo Basile SRL), and exploratory behaviour was recorded and tracked for 5 minutes using EthoVision XT14 software (Noldus Information Technology). Open arm time was chosen as a proxy measure for anxiety-like behaviours and was normalised to controls for all experiments, including analysis across time.

### 4.7 Immunohistochemistry

Mice were euthanised with an overdose of pentobarbital (Dolethal; Vetoquinol) and perfused transcardially with heparinised saline (5000IU/ml) followed by 4% paraformaldehyde (PFA). Brain and spinal cord tissue was dissected and post-fixed in 4% PFA for 2 hours, transferred to 30% sucrose and kept at 4°C until ready for sectioning. For c-fos immunostaining, free floating spinal cord sections of 40µm were blocked in 3% normal goat serum and incubated in anti-c-fos (1:5000; Synaptic Systems) for three nights. After staining slides were imaged using the Zeiss Axioscan slide scanner and cFos expressing cells were counted using Image J.

### 4.8 RT-qPCR gene expression analysis

Fresh tissue was collected following euthanasia with CO_2._ For the RS tissue experiments, mouse spinal cords were dissected into quadrants and the 2 dorsal quadrants were pooled for analysis. Cords of intra-plantar CFA injected mice were separated into ipsilateral and contralateral quadrants to site of injury and ipsilateral dorsal quadrants were processed. Total RNA was extracted using an acid phenol extraction method and processed using the RNeasy mini kit (Qiagen). 500ng of total RNA was reverse transcribed to cDNA via a two-step method, described in detail in the supplementary methods. cDNA was stored at -20°C until further processing. RT-qPCR reactions were run using the DNA Engine and the SYBR Green JumpStart Taq Ready Mix (Sigma), in the standard 3-step SYBR green heat cycle. Primer sequences for gene targets are listed in Table S3. The ratio of the relative expression of target genes to HGPRT expression was calculated using the 2ΔCt formula (Further details in supplementary methods).

### 4.9 ELISA Blood CORT levels

End of life blood was collected and processed, and circulating levels of corticosterone were measured using an ELISA kit (Abcam) as before^20^.

### 4.10 Statistical Analysis of Behavioural and Molecular Data

Statistical analyses were performed using Graph Pad Prism (Version 9.5) and SPSS IBM SPSS Statistics (Version 29). Analysis was performed with either T-tests, Analysis of Variance (ANOVA) (repeated measures when appropriate) or via Simple Linear Regression. Post hoc analysis was dependent on the test employed and performed by either Tukey’s multiple comparison test (2-way ANOVA), Holm-Sidak’s multiple comparison test (1-way ANOVA), or Sidak’s multiple comparison test (2-way ANOVA) as appropriate. Significance level was set at p<0.05. Data is presented as mean ±SEM, with or without single data points, as appropriate.

### 4.11 RNA and DNA extraction/library preps

DNA and RNA was extracted from the same lumbar spinal cord samples (L4 to L6) quadrants using the Qiagen All Prep DNA/RNA/miRNA kit as per manufacturer’s instructions. DNA and RNA were quantified using a Nanodrop 8000 spectrophotometer.

RNA integrity was measured using the Agilent 2100 Bioanalyser. All samples passed quality control with RIN scores >7. mRNA libraries were prepared from total RNA using NEBNext Ultra II Directional Library Preparation Kit. mRNA fragmentation and first strand cDNA synthesis were performed according to manufacturer’s recommendations for an insert size of 300bp (94°C for 10 minutes) and amplified for 13 cycles of PCR. Resulting libraries were quantified using the Qubit 2.0 spectrophotometer and average fragment size assessed using the Agilent 2200 Tapestation. A final sequencing pool was created using equimolar quantities of each sample library. 75bp paired-end reads were generated for each library using a NextSeq®500 in conjunction with the NextSeq®500 v2 High-output 150-cycle kit (Illumina). DNA samples were assessed for integrity using the Agilent Genomic DNA ScreenTape and reagents. Bisulphite conversion for DNAm analysis was performed using the Zymo EZ-DNA Methylation Kit using 500ng of gDNA as input.

### 4.12 RNA sequencing

FASTQ sequencing reads were assessed for quality scores (QS) and trimmed using a sliding window operation (average QS > 20). Transcripts per million of mouse mm10 reference alignment features (n = 27179) were normalised using the trimmed mean of M-values (median library size of 15.6 million counts) and filtered by expression among each group ≥10 counts. Differential gene expression was assessed by linear modelling in RStudio (>version 4.3.1) using cell type composition estimates of each RNA sample as covariates, established by querying analytically robust RNA count matrices against the mouse single-cell atlas ^59^). Bulk RNA most closely resembled 4 single cell types: myelinating oligodendrocytes, oligodendrocyte precursor cells, neurones and astrocytes (all with r > 0.6). There was no apparent difference between RS and control mice in cell types (Fig.S3). Genes exceeding a 1.2-fold change in expression and nominal p values < 0.05 were considered differentially expressed among groups.

### 4.13 DNA methylation analysis

DNA methylation was assessed using Infinium Mouse Methylation (285k) BeadChip (Illumina, USA) processed via standard protocols at the QMUL Genome Centre, Blizard Institute, Queen Mary University, London, UK. Bisulfite converted DNA was hybridised to the array and read via the Illumina iScan using the manufacturer’s standard protocol to generate red/green channel idat files. Bisulfite conversion success was confirmed prior to analysis using the mean signal intensities of CpCs over TpCs in the green channel i.e., GCT score (Green CpC to TpC) > 1^46^.

#### Data pre-processing

Probes on the sex chromosomes X and Y, and those previously identified as cross reactive,with polymorphic CpGs, or common SNVs under probe binding sites^60^, were removed. Poor samples were detected and removed if >=10% of probes possessed <3 beads or failed the detection p value threshold (>0.01). Probes with poor detection P values (>0.01 in >=10% samples), and bead counts (<3 beads per signal in >=10% samples) were eliminated. Dye and probe type biases were normalised using the subset-quantile within array method and DNA methylation status (β) calculated for all remaining probes (n=192917 and n=191134 probes in RS vs control and SAFit2 vs vehicle, respectively). Differential DNA methylation was performed using linear modelling on log2 transformed β value ratios (M-value)^61^ using estimates of cell type composition established from bulk RNA sequencing data from the same samples and surrogate variables established using SVA [PMID:17907809] as covariates. Differentially methylated probes (DMPs) with nominal significance of p < 0.05 were advanced for additional exploratory analysis using GREAT v4.0.4 ^27^. Default GREAT (Mouse: GRCm38) association parameters (basal proximal 5kb upstream and 1kb downstream, plus distal extension <=1Mb) were used with the location of the post-QC mouse array CpGs uploaded as the background regions set for comparisons. CpGs were annotated using the mouse 285k (mm10) manifest file and according to mouse neural tube (E15.5) chromatin segmentation data using the Know Your CG knowledgebase available through SeSAMe^26, 60^.

### 4.14 Intersection between DNAm and RNAseq

All cited mouse genomic co-ordinates are build GRCm38 (mm10). CpGs occurring in active promoter-like chromatin states (Tss, TssFlnk) in neural tube tissue were grouped by gene derived from the mouse 285k (mm10) manifest file and the average fold-change calculated from the differential methylation analysis. These were mapped to the average fold-change in corresponding RNA count matrices derived from the same samples^26, 60^.

### Study approval

All experiments were carried out under the Home Office License P8F6ECC28, ARRIVE guidelines were followed, and all efforts were made to minimize animal suffering and to reduce the number of animals used (UK Animal Act, 1986)

## Data availability

Data generated in this study will be made available upon completion of further analyses. Once fully processed and validated, the datasets will be accessible upon reasonable request from the corresponding authors.

### Author contributions

**Study conception and experimental designs:** OM and SMG. **Data collection:** All behavioural studies were done by OM; SMG contributed to tissue collection; TS prepared the SAFit2-VPG. EW and CAM ran the sequencing and DNA methylation arrays. **Data analysis and interpretation:** OM and SMG analysed and interpreted all data. SS ran all RNA sequencing and DNA methylome analysis guided by CGB. SH provided critical insights throughout the study. **Manuscript preparation**: OM, SS and SG wrote the initial draft of the manuscript. CGB and SH provided critical comments and revisions on the draft and all authors read and approved the final manuscript.

## Acknowledgments

The authors thank Greg Dussor, UT Dallas, for discussion on the RS model.

## Funding

his project was funded by a Brain Research UK PhD studentship to OM. SS is a member of the Advanced Pain Discovery Platform and supported by a UKRI and Versus Arthritis grant (MR/W002566/1).

## Competing interests

The authors have no competing interests related to the results or compounds in this work.

## Notes

### Competing Interest Statement

The authors have declared no competing interest.

