## Supplementary Data for "Stress, Epigenetic Remodeling and FKBP51: Pathways to Chronic Pain Vulnerability"

### Supplementary Material

#### Extended Methods

##### Drug administration

The FKBP51 inhibitor, SAFit2, was delivered as described below, and dissolved in one of the following two vehicles:

- Vehicle A: Subcutaneous injection of a slow-release vesicular phospholipid gel formulation (VPG), consisting of 50% (by mass) egg lecithin containing 80% phosphatidylcholine in 10mM phosphate-buffered saline (PBS), as before [1]
- Vehicle B: Twice daily IP injections of a solution containing 20% EtOH, 40% PPG, 5% PEG400 and 5% TWEEN 80 in 0.9% saline.

##### Elevated plus Maze (EPM)

The elevated plus maze [2,3] was used to measure anxiety-like behaviour. Mice were placed on the centre of the EPM, which consisted of two open arms (35cm) and two closed arms (35cm), joined by a centre section (5cm x 5cm) and elevated 60cm from the ground (Ugo Basile SRL). Exploratory behaviour of the open and closed arms was recorded for 5 minutes. The maze was wiped clean with 70% ethanol between mice. EthoVision XT14 (Noldus Information Technology) allowed for tail-nose tracking of total time spent in each arm, as well as distance travelled. Open arm time was selected as a proxy measure for anxiety-like behaviours. Any mice who fell from the maze during testing were excluded from the analysis. Time spent in the open arms was normalised to controls for all experiments, including analysis across time.

##### Open Field (OF)

The open field was used to measure locomotion. Mice were placed in the centre of a round arena (30cm diameter) and distance travelled in 5 minutes was recorded. EthoVision XT14 (Noldus Information Technology) was used to track centre points of mice.

##### Immunohistochemistry

Mice were anaesthetised in an anaesthetic chamber (2% isoflurane in oxygen, delivered at 2L/min) until unconscious and euthanised with an overdose of pentobarbital (Dolethal; Vetoquinol). They were then perfused transcardially with heparinised saline (5000IU/ml), followed by 4% paraformaldehyde (PFA) in 0.1M phosphate buffer (PB). The brain and spinal cords were dissected and post-fixed in 4% PFA for 2 hours, transferred to 30% sucrose and kept at 4°C until ready for sectioning.

For c-fos immunostaining, free floating spinal cord sections of 40um thickness were first blocked in 3% normal goat serum, 3% Triton X and 2% hydrogen peroxide in 0.1M PB for 1 hour. Sections were then incubated in anti-cFos (1:5000; Synaptic Systems) diluted in Tris-buffered saline and tween 20 (TTBS), for three nights at 4C. The primary antibody was removed, and the sections were incubated in biotinylated secondary antibody for 2 hours (1:500; Vector), followed by an avidin-biotin complex (ABC: 1 hour; Vector). Finally, the stain was developed using a DAB kit (Vector) for up to 10 minutes. Sections were then mounted and left to dry overnight, before dehydrating in increasing concentrations of ethanol, and coverslipped with DPX.

##### RT-qPCR gene expression analysis

Fresh tissue was collected following euthanasia with $CO_2$. For the RS tissue experiments, mouse spinal cords were dissected into quadrants and the 2 dorsal quadrants (i.e. both left and right side) were pooled for analysis. Cords of intra-plantar CFA injected mice were separated into ipsilateral and contralateral quadrants to site of injury and ipsilateral quadrants were processed. Total RNA was extracted using an acid phenol extraction method (TRIzol reagent; Qiagen), which involved homogenisation by hand and passage through a biopolymer-shredder (QIAshredder; Qiagen). The RNeasy mini kit (Qiagen) was used to complete the extraction and RNA concentrations were measured using a Nanodrop. 500ng of total RNA was then reverse transcribed to cDNA via a two-step method. First, samples were incubated with random nonamers (2uM; Sigma), oligo $dT_{20}$ primer mix (1uM; Promega) and 10mM dNTP mix (0.5uM; Promega) at 65°C for 5 minutes. First-strand buffer (5x) and superscript III were then added, along with DTT (0.1M; all Promega), RNaseIN recombinant ribonuclease inhibitor (Promega) and completed by exposure to a heat cycle of 25°C for 5 minutes, 50°C for 50 minutes and 70°C for 15 minutes. Appropriate positive and negative controls were included in the process. cDNA was stored at -20°C until further processing. RT-qPCR reactions were run using the DNA Engine and the SYBR Green JumpStart Taq Ready Mix (Sigma), in the standard 3-step SYBR green heat cycle. Primer sequences for gene targets are listed in Table S1. Reactions were run in triplicates and assessed by melting curve analysis. The ratio of the relative expression of target genes to HGPRT expression was calculated using the 2ΔCt formula.

**ELISA Blood CORT levels**

End of life blood was collected by syringe and placed in an Eppendorf containing 4% sodium citrate (1:9 ratio to blood). Tubes were then placed on ice for two hours and centrifuged at 10000 rpm for 15 minutes at 4°C. Serum was transferred to a clean tube. Circulating corticosterone levels were measured using an ELISA kit (Abcam).

**RNA and DNA extraction/library preps**

DNA and RNA was extracted from the same spinal cord samples lumbar (L4 to L6) quadrants using the Qiagen AllPrep DNA/RNA/miRNA kit as per manufacturer's instructions. DNA and RNA were quantified using a Nanodrop 8000 spectrophotometer.

RNA integrity was measured using the Agilent 2100 Bioanalyser. All samples passed Quality Control (QC) with RIN scores >7. mRNA libraries were prepared from total RNA using NEBNext Ultra II Directional Library Preparation Kit. Fragmentation of isolated mRNA prior to first strand cDNA synthesis was carried out using incubation conditions recommended by the manufacturer for an insert size of 300bp (94°C for 10 minutes). 13 cycles of PCR were performed for final library amplification. Resulting libraries were quantified using the Qubit 2.0 spectrophotometer and average fragment size assessed using the Agilent 2200 Tapestation. A final sequencing pool was created using equimolar quantities of each sample library. 75bp paired-end reads were generated for each library using the Illumina NextSeq®500 in conjunction with the NextSeq®500 v2 High-output 150-cycle kit to obtain an average of 19.5M read pairs per sample.

DNA samples were assessed for integrity using the Agilent Genomic DNA ScreenTape and reagents. Bisulphite conversion was performed using the Zymo EZ-DNA Methylation Kit using 500ng of gDNA as input.

**RNA sequencing**

RNA sequencing data were pre-processed and analysed as previously described [4]. Briefly, raw paired-end next generation sequencing reads (Illumina HiSeq 2500) were assessed for quality scores and trimmed using a sliding window operation with an average quality score $Q > 20$. Transcripts per million of mouse mm10 reference alignment features ($n = 27179$) were normalised using the trimmed mean of M-values (median library size of 15.6 million counts) and filtered by expression (≥10 counts) to remove genes that are either not expressed or lowly expressed among all samples ($n = 11646$ genes). In summary, 15533 unique genes were advanced to downstream analysis.

Differential gene expression was established using linear modelling in RStudio (>version 4.3.1). Bulk RNA count matrices were queried against the single-cell mouse cell atlas (Han et al., 2018) to estimate cell type composition. Bulk RNA samples most closely resembled 4 single cell types: myelinating oligodendrocytes, oligodendrocyte precursor cells, neurones and astrocytes (all with $r > 0.6$). There was no apparent difference between RS and control mice in cell types (Fig.S5). Cell type resemblance z-scores estimated by the mouse cell atlas were included as covariates in differential expression (RS – Control). Genes exceeding a 1.2-fold change in expression and nominal p values < 0.05 were considered differentially expressed among groups.

**DNA methylation analysis**

DNA methylation was assessed using Infinium Mouse Methylation (285k) BeadChip (Illumina, USA) processed via standard protocols at the QMUL Genome Centre, Blizard Institute, Queen Mary University, London, UK. Bisulfite converted DNA was hybridised to the array and read via the Illumina iScan using the manufacturer's standard protocol to generate red/green channel idat files. Bisulfite conversion was assessed using the mean signal intensities of CpGs over TpGs in the green channel. Conversion success scores (± SD) of 1.12 ± 0.010 and 1.12 ± 0.014 in control and RS mice, respectively, reveal similar and complete bisulfite conversion between groups.

*Data pre-processing*

Probes on the sex chromosomes X and Y, and those previously identified as poorly reactive [4] were removed. Poor samples were detected and removed if >=10% of probes possessed <3 beads or failed the detection p value threshold (>0.01). Probes with poor detection P values (>0.01 in >=10% samples), and bead counts (<3 beads per signal in >=10% samples) were eliminated. Dye and probe type (I or II) biases were normalised using the subset-quantile within array method and DNA methylation status (as β values) were calculated for all remaining probes (n = 192917 and n = 191134 probes in RS vs control and SAFit2 vs vehicle, respectively). Differential DNA methylation was performed on log2 transformed β value ratios (M-value) using the contrast RS – Control or SAFit2- Vehicle. Estimates of cell type composition established using bulk RNA sequencing data from the same samples were included as covariates in the linear model along with surrogate variables established using SVA. Differentially methylated probes (DMPs) with nominal significance of p < 0.05 were advanced for additional exploratory analysis including genomic enrichment of cis-regulatory regions using GREAT [5]. CpGs were annotated using the mouse 285k (mm10) manifest file and visualised using the UCSC genome browser.

**Intersection between DNAm and RNAseq.**

CpGs were annotated according to mouse neural tube (E15.5) chromatin segmentation data using the Know Your CG knowledgebase available through SeSAMe [4,6]. CpGs occurring in active promoter-like chromatin states (Tss, TssFlnk) were grouped by gene derived from the mouse 285k (mm10) manifest file and the average fold-change calculated from the differential methylation analysis. These were mapped to the average fold-change in corresponding RNA count matrices derived from the same samples.

#### Supplementary Data

**Table S1: Gene Set enrichment via GREAT of loci modulated by RS.** Significant (nominal $p < 0.05$) CpGs ($n = 11644$) were tested for gene set enrichment via GREAT using analytically robust mm10 array probes ($n = 192917$) as the background region. Distal gene regulatory domains were extended up to 1000 kb from the TSS.

| Term | | Fold enrichment | FDR adjusted p value |
|---|---|---|---|
| **Biological process** | Ubiquitin homeostasis | 8.28 | 0.0238 |
| | pre-miRNA export from nucleus | 13.25 | 0.0344 |
| | Positive regulation of RNA interference | 13.25 | 0.0344 |
| | Aspartate catabolic process | 3.11 | 0.0372 |
| | Glutamate catabolic process to aspartate | 3.18 | 0.0419 |
| | Glutamate catabolic process to 2-oxoglutarate | 3.18 | 0.0419 |
| | Glutamate catabolic process | 2.82 | 0.0494 |
| | Regulation of protein localisation to ciliary membrane | 8.28 | 0.0486 |
| | Retrograde axonal transport | 2.31 | 0.0478 |
| **Molecular function** | Heterotrimeric G-protein binding | 3.93 | 0.007 |
| | pre-miRNA transporter activity | 13.25 | 0.0373 |
| | L-phenylalanine aminotransferase activity | 2.98 | 0.0328 |

**Table S2: *Fkbp5* associated probes in both RS *vs* Control and SAFit2 *vs* Vehicle studies**. CpGs are mapped to their closest gene, distance and chromatin state using mouse neural tube (E15.5) segmentation. FC: log2-fold change contrast. HMM: Chromatin state signatures established using hidden Markov model. Chr: Chromosome

() = associated by GREAT

[b] = bases

Note: cg34792474_TC21 falls in Tss of neighbouring gene *Armc12* (distance to Tss is +284). Predicted by GREAT to interact with Fkbp5 (+6031). The distance of this CpG to the Tss defined by cg34791938_TC11 (chr17:28,486,150) is predicted as -37427

| Probe ID | Location | chrHMM | CpG Island | Distance to TSS [*b*] | $Log_2FC$ [RS v C] | p [RS v C] | $Log_2FC$ [SAF v V] | p [SAF v V] |
|---|---|---|---|---|---|---|---|---|
| cg34791125_BC21 | chr17:28419801 | EnhPois | OpenSea | 8829 | 0.10 | 0.402 | -0.18 | 0.377 |
| cg34791127_TC21 | chr17:28419929 | EnhPois | OpenSea | 8701 | -0.09 | 0.374 | 0.18 | 0.196 |
| cg34791132_BC11 | chr17:28420600 | EnhPois | OpenSea | 8030 | -0.06 | 0.527 | -0.34 | 0.010 |
| cg34791341_BC21 | chr17:28441672 | QuiesG | OpenSea | -618 | -0.08 | 0.301 | 0.12 | 0.316 |
| cg34791429_BC21 | chr17:28448556 | QuiesG | OpenSea | 37594 | -0.02 | 0.810 | 0.05 | 0.613 |
| cg34791585_TC21 | chr17:28466852 | - | OpenSea | 19298 | 0.02 | 0.893 | -0.08 | 0.496 |
| cg34791595_BC11 | chr17:28467845 | TssFlnk | OpenSea | 18305 | -0.03 | 0.787 | -0.06 | 0.687 |
| cg34791627_TC21 | chr17:28470968 | EnhPois | OpenSea | 15182 | -0.10 | 0.253 | -0.11 | 0.389 |
| cg34791655_BC21 | chr17:28473058 | QuiesG | OpenSea | 13092 | 0.01 | 0.931 | -0.09 | 0.542 |
| cg34791666_TC21 | chr17:28474245 | QuiesG | OpenSea | 11905 | -0.06 | 0.687 | -0.01 | 0.966 |
| cg34791916_BC21 | chr17:28486157 | Tss | S_Shore | -7 | -0.07 | 0.441 | -0.08 | 0.579 |
| cg34791938_TC11 | chr17:28486463 | Tss | S_Shore | -313 | -0.24 | 0.043 | -0.03 | 0.799 |
| cg34791125_BC21 | chr17:28419801 | EnhPois | OpenSea | 8829 | 0.10 | 0.402 | -0.18 | 0.377 |
| cg34792474_TC21 | chr17: 28523577 | Tss | S_Shore | (-37427) | -0.22 | 0.032 | 0.18 | 0.196 |

**Table S3: Primers sequences for RTqPCR.**

| Target gene | Forward Sequence (5'-3') | Reverse Sequence (5'-3') |
|---|---|---|
| ***Fkbp5*** | CAATGCTGAGCTTATGTACG | CTTTTCTTGGTGTCCATCTC |
| ***Nr3c1*** | CTGGACGGAGGAGAACTCAC | GGACAACCTGACTTCCTTGG |
| ***Grα*** | GAGCACACCAGGCAGAGTTT | AGGCCGCTCAGTGTTTTCTA |
| ***Grβ*** | CCATAATGGCATACCGAAGC | AGGCCGCTCAGTGTTTTCTA |
| ***HPRT*** | AGGGATTTGAATCACGTTTG | TTTACTGGCAACATCAACAG |

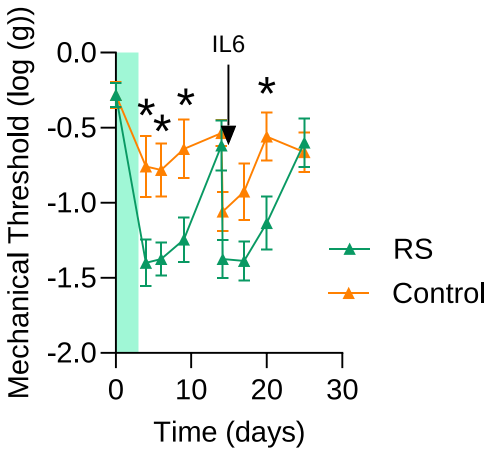


**Figure S1: Sub-chronic stress primes for hyper-responsiveness to IL6-induced inflammatory pain.** Hind paw mechanical withdrawal thresholds were measured in male mice. RS-induced a significant reduction in hind paw thresholds compared to control and exacerbated the response to intra plantar IL6 injection. IL6 was injected on day 14 (25µl of 0.1ng IL6). N=8/8. RM ANOVA, day 0 to day 14: $F_{1,14}$=13.1, p=0.003; day 14 +6h to day 20: $F_{1,14}$=7.4, p=0.017. *P<0.05: post-hoc analysis one-way ANOVA. Green panel indicates the 3 days of RS paradigm.

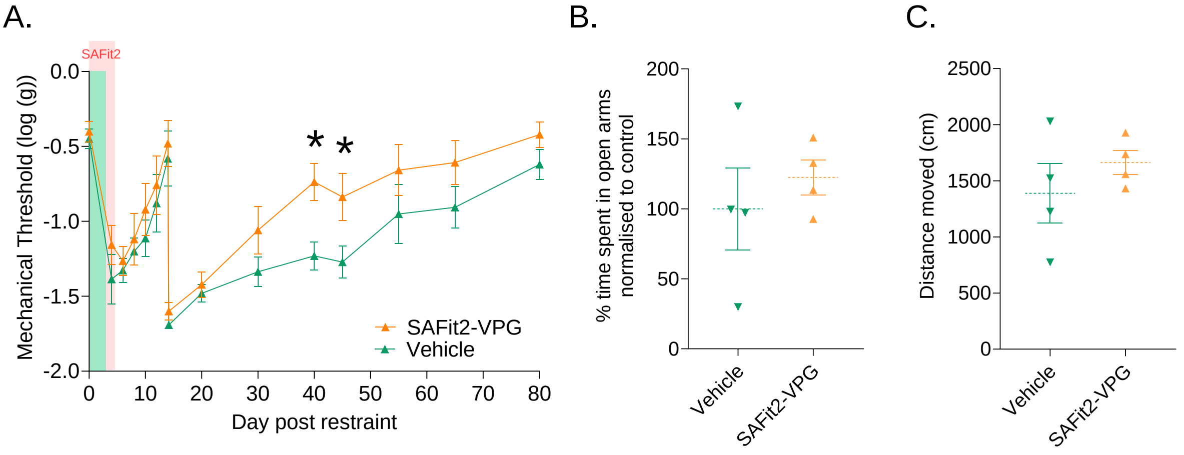


**Figure S2: Inhibition of FKBP51 using repeated i.p. injection of SAFit2 reduces CFA induced mechanical hypersensitivity.** (**A**) Hind paw mechanical withdrawal thresholds were measured using Von Frey filaments in male mice receiving either Vehicle or SAFit2 injections, administered twice daily by i.p. injections for 5 days from two days prior to the RS paradigm. All mice received intra plantar CFA was on day 14. SAFit2 prevented the CFA-induced exacerbation of CFA-induced hypersensitivity. *P<0.05, Univariate ANOVA at 45 days and 60 days. RM ANOVA; Time 14 days + 6h to 80 days: $F_{1,10}$=6.1, P=0.033. (**B, C**) Control experiments involved the subcutaneous administration of SAFit2- or Vehicle-VPG (control) to naive mice. Percentage of time spent exploring the open arms of the EPM (B) and distance travelled in the Open Field (B) was measured 5 days after receiving the injection. The SAFit2-VPG injected mice were not significantly different from Vehicle in either measure.

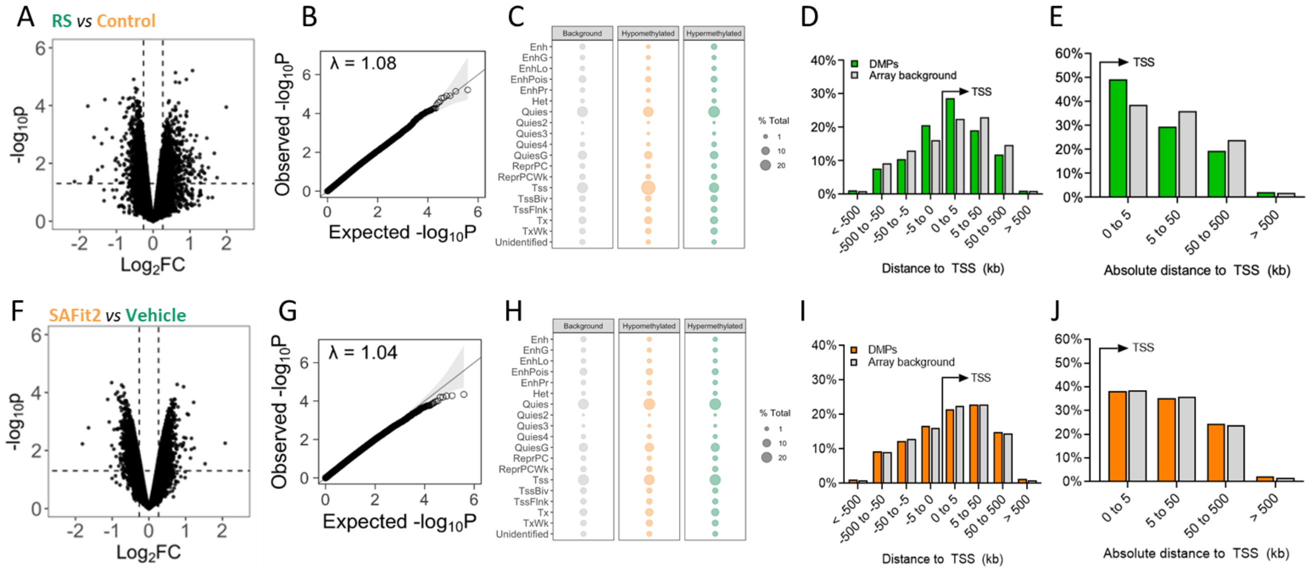


**Figure S3: Summary of differentially methylated probes (DMPs): RS *vs* Control and SAFit2 *vs* Vehicle.** (**A, F**) Volcano Plot for the differential methylation results from DNA methylation analysis comparing RS versus control mice (A) and SAFit2 vs Vehicle (F). Each point on the plot represents a single CpG. The x-axis denotes the log2 fold change (FC) in methylation between the two conditions, with positive values indicating increased methylation in the RS and RS + SAFit2 groups and negative values indicating increased methylation in Control and RS + vehicle groups. The y-axis represents the -log10 transformed p-value, indicating the statistical significance of the differential methylation. (**B, G**) Quantile–quantile plot showing the distribution of observed -log10 p values from differential methylation analysis vs their expected distribution under the null hypothesis and genomic inflation factor (λ). (**C, H**) CpGs with nominal ($p < 0.05$) significance were mapped to genomic location and 15 state mouse neural tube chromatin segmentation data (E15.5) using the KYCG database (see Methods). Background represents all CpGs remaining on the beadchip following quality control assessments. Data are scaled to a percentage of total CpGs in each comparison. (**D, E, I, J**): GREAT genomic enrichment of nominal CpGs binned by orientation (**D, I**) and absolute (**E, J**) distance to the nearest transcription start site (TSS). CpGs identified as significant ($p < 0.05$) in the RS vs control contrast (A) were enriched compared to background CpGs within 5kb of the nearest TSS.

A.

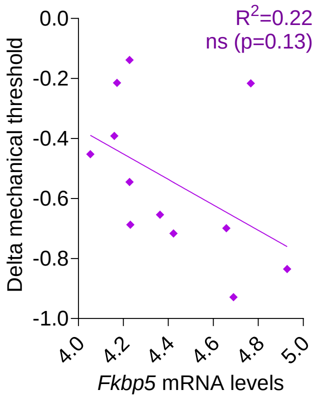


**Figure S4: There is no correlation between *Fkbp5* mRNA levels and mechanical thresholds.** *Fkbp5* mRNA levels were assessed by sequencing.

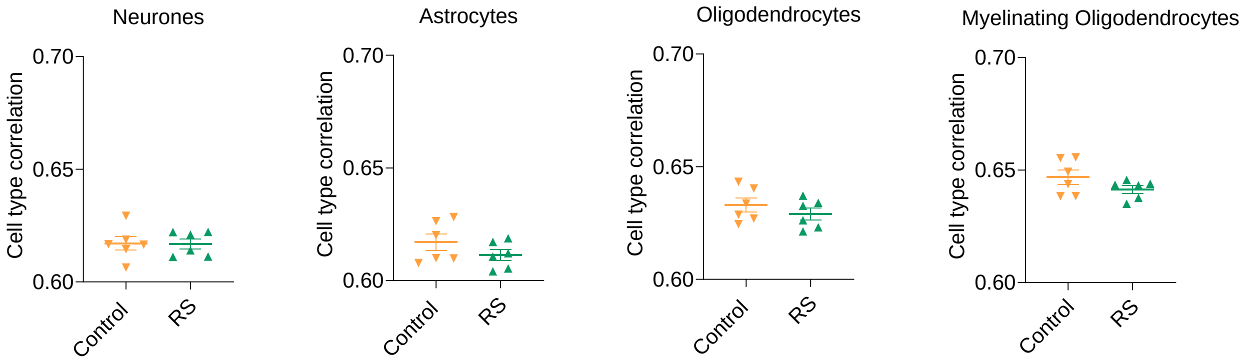


**Fig.S5: Correlation analysis between the major cell types and spinal cord samples showed no difference between RS and control mice.** Correlation coefficients between spinal cord samples and the predominant cell types expressed in spinal cord tissue were evaluated using RNA sequencing data. The degree of correlation with neuronal cell type, astrocytes, oligodendrocytes and myelinating oligodendrocytes was no different between RS and control mice spinal cord samples. These results suggested that cell type proportions had not changed after RS.
